# The Beacon Calculus: A formal method for the flexible and concise modelling of biological systems

**DOI:** 10.1101/579029

**Authors:** Michael A. Boemo, Luca Cardelli, Conrad A. Nieduszynski

## Abstract

Biological systems are made up of components that change their actions (and interactions) over time and coordinate with other components nearby. Together with a large state space, the complexity of this behaviour can make it difficult to create concise mathematical models that can be easily extended or modified. This paper introduces the Beacon Calculus, a process algebra designed to simplify the task of modelling interacting biological components. Its breadth is demonstrated by creating models of DNA replication dynamics, the gene expression dynamics in response to DNA methylation damage, and a multisite phosphorylation switch. The flexibility of these models is shown by adapting the DNA replication model to further include two topics of interest from the literature: cooperative origin firing and replication fork barriers. The Beacon Calculus is supported with the open-source simulator bcs (https://github.com/MBoemo/bcs.git) to allow users to develop and simulate their own models.

**Author summary:** Simulating a model of a biological system can suggest ideas for future experiments and help ensure that conclusions about a mechanism are consistent with data. The Beacon Calculus is a new language that makes modelling simple by allowing users to simulate a biological system in only a few lines of code. This simplicity is critical as it allows users the freedom to come up with new ideas and rapidly test them. Models written in the Beacon Calculus are also easy to modify and extend, allowing users to add new features to the model or incorporate it into a larger biological system. We demonstrate the breadth of applications in this paper by applying the Beacon Calculus to DNA replication and DNA damage repair, both of which have implications for genome stability and cancer. We also apply it to multisite phosphorylation, which is important for cellular signalling. To enable users to create their own models, we created the open-source Beacon Calculus simulator bcs (https://github.com/MBoemo/bcs.git) which is easy to install and is well-supported by documentation and examples.

## Introduction

The ability to quickly create flexible and concise models of biological systems makes mathematical modelling more practical, enables rapid hypothesis testing, and increases the likelihood that modelling will be used to ensure that conclusions drawn from experiments are consistent with data. Process calculi (or process algebras) are valuable tools for assessing the performance, reliability, and behaviour of a system. Each component in a system is abstracted as a process that can perform actions. Communication actions allow processes to interact with one another to perform coordinated behaviours. The semantics of a process calculus sets rigorous rules that govern which actions that processes can perform, enabling formal reasoning about whether a system is ever capable of performing (or not performing) a certain sequence of actions. While process calculi have been historically developed to formally reason about programs and algorithms, they are applicable to any concurrent system (such as biological systems).

There have been many process calculi developed in recent decades: The calculus of communicating systems (CCS) [1] and communicating sequential processes (CSP) [2] are early and foundational examples of process calculi where “reachability”, or whether a system can ever perform a certain set of actions, can be determined using the language’s structural operational semantics. Performance Evaluation Process Algebra (PEPA) assigned a rate to each action so that the system could be mapped onto a continuous time Markov chain (CTMC) [3, 4]. Once expressed as a CTMC, the system can be simulated by generating random paths through the CTMC’s states. It also becomes possible to determine the probability that a behaviour occurs within a specified amount of time, and the system’s asymptotic behaviour can be determined using the CTMC’s stationary distribution [5]. Tools have been developed to map the PEPA language onto a CTMC and perform this analysis, including the PEPA workbench [6], the PEPA Eclipse plug-in [7], and a PEPA-to-PRISM compiler [8].

PEPA has been expanded in a number of ways: Bio-PEPA [9] is an extension for the simulation and verification of biochemical networks and can be analysed via the accompanying Eclipse plugin or the Bio-PEPA workbench [10]. PEPAk is an extension of PEPA that includes process parameters and gated actions [11, 12]. PEPA has also been used as an inscription language for stochastic Petri nets, providing a natural framework for modelling mobile systems [13].

The *π*-calculus encodes models of concurrent processes using a notion of naming, whereby processes can use channels to communicate channel names to dynamically change which processes can communicate with one another [14]. The stochastic *π*-calculus is an extension of the *π*-calculus that has been used for performance modelling in a number of biological applications [15]. SPiM is a stochastic *π*-calculus simulator for large numbers of interacting biological molecules [16]. In addition, several studies have use the stochastic *π*-calculus to model regulatory networks in biology, for example [17–20].

This paper introduces the Beacon Calculus, which makes it simple and concise to encode models of complex biological systems. It is a tool that builds upon the intuitive syntax of PEPA and mobility in the *π*-calculus to produce models that are shorter, simpler, and more flexible than they would be if they were encoded in either of these languages (Section S2). The following section gives a description of the language by way of examples (a formal description of the language is given in Section S1). To demonstrate breadth, results are presented for Beacon Calculus models of three different biological systems from the literature, each of which highlights one of the language’s main features: a model of DNA replication dynamics that fits replication timing data, a model of the gene expression response to DNA methylation damage in which the model qualitatively matches single-cell tracking experiments, and a stochastic version of an established deterministic multisite phosphorylation model from the literature.

## Results

This section begins with an introduction to the Beacon Calculus by way of examples. Usage is demonstrated by gradually building upon a simple model of a bimolecular reaction *A* + *B* ↔ *AB*, leading to a complex yet concise model that uses many of the language’s features. (An additional introductory example describing kinesin stepping down a microtubule is provided in Section S5.) In addition to the Beacon Calculus itself, a contribution of this paper is bcs, an open-source Beacon Calculus simulator (https://github.com/MBoemo/bcs.git). To make it clear how to translate theory to practice, all examples are given in bcs source code so that they can be simulated and experimented with. A more formal and precise specification of the language and its semantics is given in Section S1. Following an outline of the language, the Beacon Calculus is then applied to three diverse areas of biological research: DNA replication, DNA damage response, and multisite phosphorylation.

### Language Overview

Models are written in the Beacon Calculus by representing components in a system as processes that can perform actions. Processes can make an exclusive choice between multiple actions, execute multiple actions in parallel, and perform actions in a sequence. These three simple but powerful combinators are common amongst many process algebras and are used in CCS, PEPA, the *π*-calculus, and others [22]. The Beacon Calculus is a stochastic process calculus where each action is specified as an ordered pair together with the rate at which it is performed. The ordered pairs `{a,ra}` and `{b,rb}` specify rates for actions `a` and `b`, respectively. The following three examples of process definitions show how each combinator is used:

- `P = {a,ra}.{b,rb}` uses the unary prefix operator “`{a,ra}`. —” to denote a sequence of actions whereby action `a` is performed at rate `ra` and, once it has finished, action `b` is performed at rate `rb`.
- `P = {a,ra} + {b,rb}` uses the choice operator “+” to denote the exclusive choice between performing action `a` and rate `ra` and performing action `b` at rate `rb`. The probability of choosing action a is 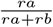, and the probability of choosing action `b` is 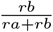.
- `P = {a,ra} || {b,rb}` uses the parallel operator “`||`” to denote that actions `a` and `b` are performed in parallel at their respective rates.

Prefix binds stronger than choice, and choice binds stronger than parallel execution. For example, in the following process

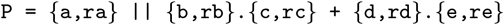

process `P` makes an exclusive choice between performing action `b` at rate `rb` and performing action `d` at rate `rd`. If `b` is chosen, `P` then performs action `c` at rate `rc` while if action `d` is chosen, `P` performs action `e` at rate `re`. All the while, `P` can perform action `a` at rate `ra` in parallel.

A process can have a finite sequence of parameters which, in practice, is often used to encode the process’s location, a quantity, or a state (though there are many other uses as well). A process `P` with parameters `i1,i2,…,in` is denoted using the notation `P[i1,i2,…,in]`. Processes can change their parameters through recursion. This is often used when a process moves (if the parameter models a location), modifies how much of something it has (if the parameter models a quantity), or otherwise changes state in some way that should influence the process’s later behaviour. For example, the following model describes a process that successively increments `i` by one and doubles `j`:

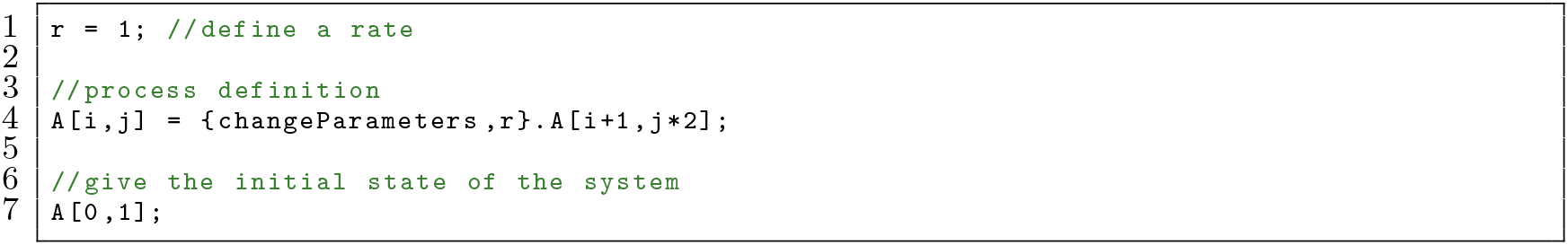

In this model, process `A` has the two parameters `i` and `j`. The system begins with one copy of `A` with values `i=0` and `j=1` (Line 7). Each time `A` performs the action `changeParameters` at rate `r`, the value of `i` is increased by one and the value of `j` is doubled.

If this model were run in bcs, `A` would continue changing the values of `i` and `j` until it hit the maximum number of transitions allowed by the software. To create effective models, it is often necessary to specify that a process should only perform an action if the parameter values meet a certain condition. A process can change its behaviour according to its parameter values by using a gate, which is a condition that must be satisfied for a process to perform an action. Gated actions are written using the notation

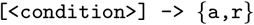

where action `a` can only be performed if the condition is true. The Beacon Calculus simulator supports the following operators in the expression for the gate condition:

- <=, less than or equal to,
- <, less than,
- >=, greater than or equal to,
- >, greater than,
- ==, equal to,
- !=, not equal to,
- &, logical and,
- |, logical or,
- ~, logical not.

For example, suppose `A` should continue while `i<5` and `j<10`. This can be expressed as follows:

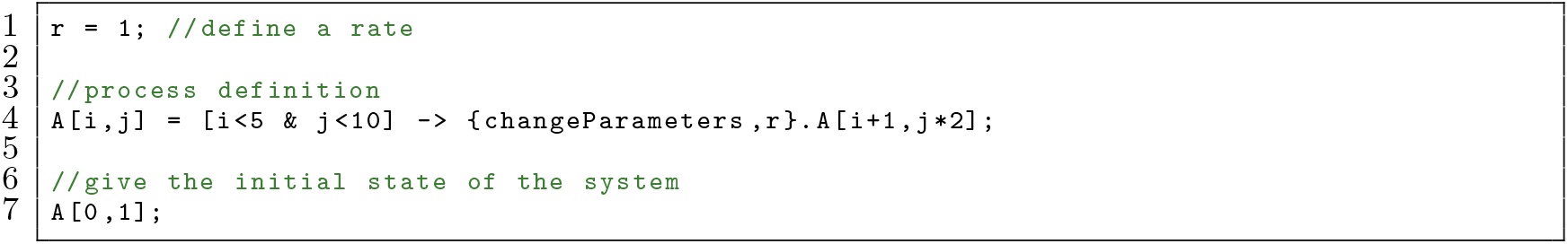

Once the condition specified in the gate no longer holds, `A` can no longer perform the action `changeParameters`. When a process can no longer perform any actions, it is said to be deadlocked and is removed from the system. If all processes in the system are deadlocked, the simulation stops. In this case, the simulation will stop when `A` has parameter values `i=4` and `j=16`.

In order for the Beacon Calculus to be useful for biological applications, a process must be able to react in some way to the actions of other processes; they must be able to communicate with one another via special actions. Handshakes are a common type of synchronous communication in many process algebras whereby two separate processes each perform an action at the same time. In the Beacon Calculus, two processes handshake when the following two actions are performed together:

- A handshake send is written {@c![i],rs}; it denotes a handshake that is offered on channel `c` that transmits parameter `i`.
- A handshake receive is written {@c?[Ω](x),rr}; it denotes a handshake that can be received on channel `c` so long as the parameter from the sending handshake is a member of the set Ω. The particular parameter received is bound to the variable `x` and can be used subsequently by the process.

A handshake always occurs between exactly two processes at a rate equal to the product of the handshake receive rate and the handshake send rate. A handshake send and a handshake receive must always be performed together. If a process is ready to send a handshake but there is no other process that can receive the handshake, then the first process must wait until another process is ready to perform the handshake receive. There is no crosstalk between channels, meaning two processes cannot handshake by performing actions {@c![i],rs} and {@d?[Ω](x),rr} because the channel names do not match. The following example shows how two reactants `A` and `B` undergo one-dimensional diffusion where they can react via a handshake when they are in the same position:

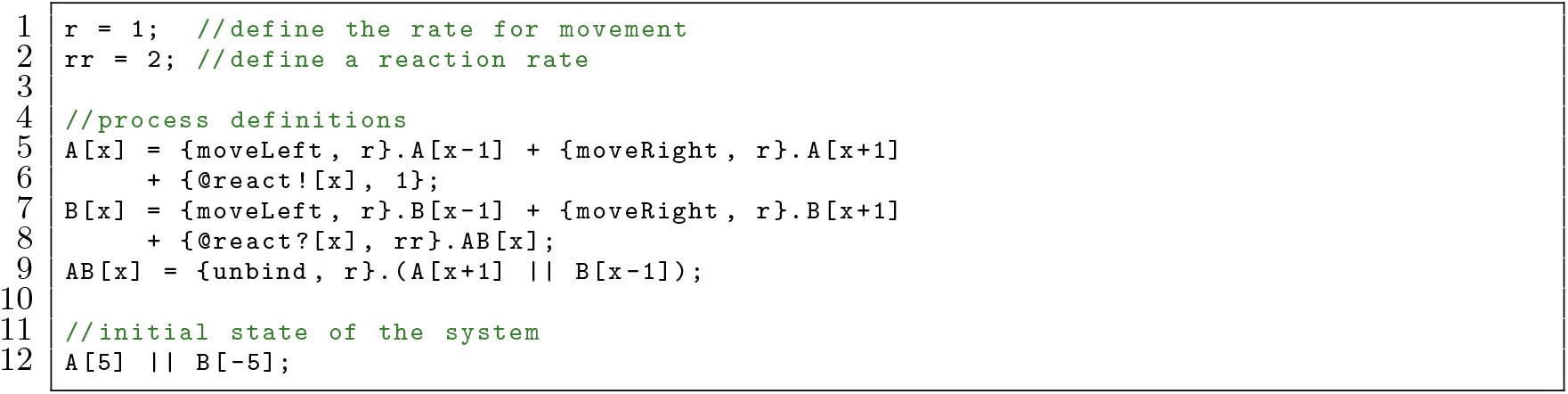

This model has two reactants, `A` and `B`, undergoing one-dimensional diffusion. `A` starts at position `i=5` and `B` starts at `i=−5` (Line 12). Both processes make a choice between stepping left at rate `r` or stepping right at rate `r` (Line 5,7). The rates are equal so the diffusion is unbiased, but biased diffusion could be introduced by making the rate for one direction higher than the other. When both `A` and `B` are at the same position, their parameters match and a handshake is possible over channel `react` at rate `rr*1=rr` (Lines 6,8). The probability of the handshake is 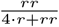. If the handshake is chosen, `A` and `B` react to form `AB` (Line 8). Once formed, `AB` unbinds to reform `A` and `B` at rate `r` (Line 9).

In the previous model, some of the code is redundant: processes `A` and `B` behave similarly, yet the `moveLeft and moveRight` actions are typed out in each case. The code can be made more concise by using parameters so that there is a reactant process `R` at position given by parameter `x` with an identity encoded by parameter `i`. `P`rocess `A` becomes reactant `R` with `i=0` and `B` becomes reactant `R` with `i=1`. This can be expressed as follows, which is equivalent to the previous model:

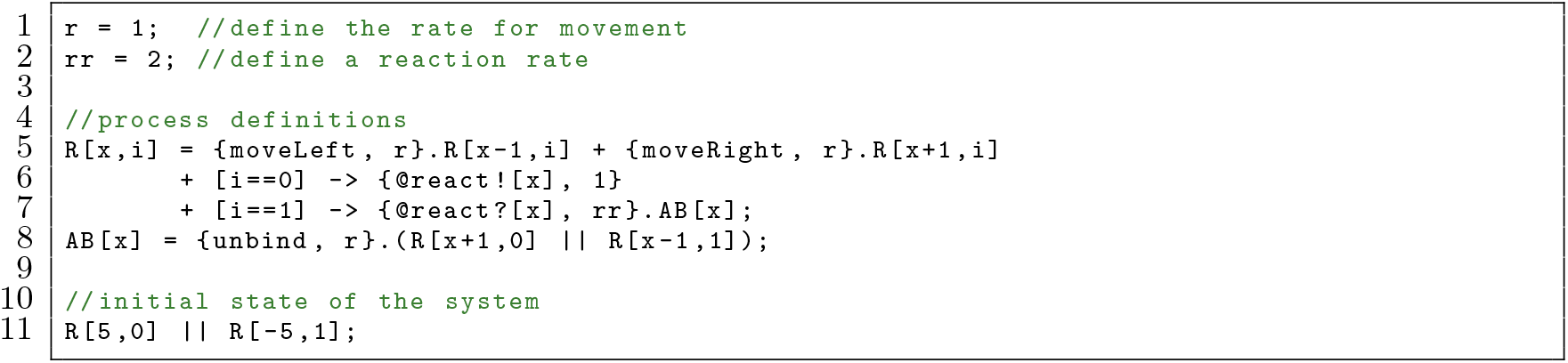

Here, the reactant `R` undergoes one-dimensional diffusion (Line 5). If it has parameter `i=0` (Line 6) then it can react with a reactant that has parameter `i=1` to form `AB` (Line 7).

While the handshake receive in the previous example could only receive a single value, handshake receives can accept a set of possible values. A set is specified in the Beacon Calculus simulator using the following operations. Examples are written for each to show the set (right) encoded by each Beacon Calculus expression (left). Note that set subtraction binds more strongly than set intersection, which in turn binds more strongly than set union.

- ‥, range,

– `0‥3 ≡` {0, 1, 2, 3}
– `−1‥2 ≡` {−1, 0, 1, 2}
- `U`, set union,

– `0‥3 U 6‥7 ≡` {0, 1, 2, 3, 6, 7}
– `−1 U 0‥3 ≡` {−1, 0, 1, 2, 3}
- `I`, set intersection,

– `0‥10 I 8‥15 ≡` {8, 9, 10}
– `0‥2 U 8‥15 I 4‥9 ≡` {0, 1, 2, 8, 9}
- \, set subtraction.

– `0‥5\3 ≡` {0, 1, 2, 4, 5}
– `0‥5\8 ≡` {0, 1, 2, 3, 4, 5}

If a handshake receive can accept multiple values, the receiving process can bind the value it receives to a variable for later use. The process may, for instance, use this value in a rate expression or as a parameter. The binding variable can be used in the rate expression to indicate how different values can be received at different rates; it can bias which value in the set is received. For example, suppose it is more likely that two kinesin motors impede each other as they get closer to one another. The two definitions for kinesin below, `B1` and `B2`, are equivalent.

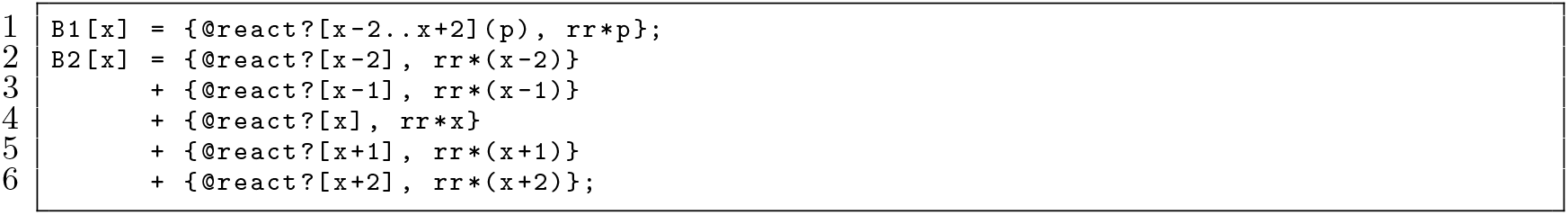

While handshakes allow two processes to perform a coordinated action simultaneously, beacons provide the means for asynchronous communication. In practice, beacons can be used to communicate the state change of a process globally to all other processes in the system. Using beacons, a process can efficiently indicate to a number of other processes that a task has been accomplished (shown in the following example) or keep track of tasks that have been done over time (shown in the DNA replication example to follow). A process can launch a beacon on a channel that transmits a parameter; the beacon stays active until it is explicitly killed by a process. While active, the beacon can be received any number of times by any process (including the process that launched it). Processes can also check whether a particular beacon is active and only carry on if there is no active beacon with a given channel and parameter.

- A beacon launch is written `{c![i], rs}`; it denotes a beacon that is launched on channel `c` that transmits parameter `i`.
- A beacon kill is written `{c#[i], rs}`; it denotes an action that kills a beacon on channel `c` transmitting parameter `i` if one exists. If one does not exist, the action is still performed but the set of active beacons does not change. Once a beacon is killed, it can no longer be received unless it is re-launched by a process.
- A beacon receive is written `{c?[Ω](x), rs}`; it denotes an action that can only be performed if there is an active beacon on channel `c` transmitting a parameter `i` in Ω. The parameter received is bound to `x` and can be used subsequently in the process.
- A beacon check is written `{~c?[Ω], rs}`; it denotes an action that can only be performed if there is no active beacon on channel `c` transmitting a parameter in Ω.

In the following example, a “clock” process `C` changes between two states, 1 and 2, at rate `rs`. When the process changes state, it launches a beacon on channel `state` with the value corresponding to the new state (Line 10). The unbinding rate of `AB` depends on the value of the parameter transmitted by the beacon: process `AB` uses the range operator to receive a value of either 1 or 2 on channel `state` and binds that value to `s` (Line 11). The value of `s` is used in the rate of the beacon receive so that if `C` is in state 1, `AB` dissociates at rate `r*1`. Likewise, `AB` dissociates at rate `r*2` if `C` is in state 2. This allows `C` to autonomously change its state and, in doing so, easily affect the behaviour of other processes.

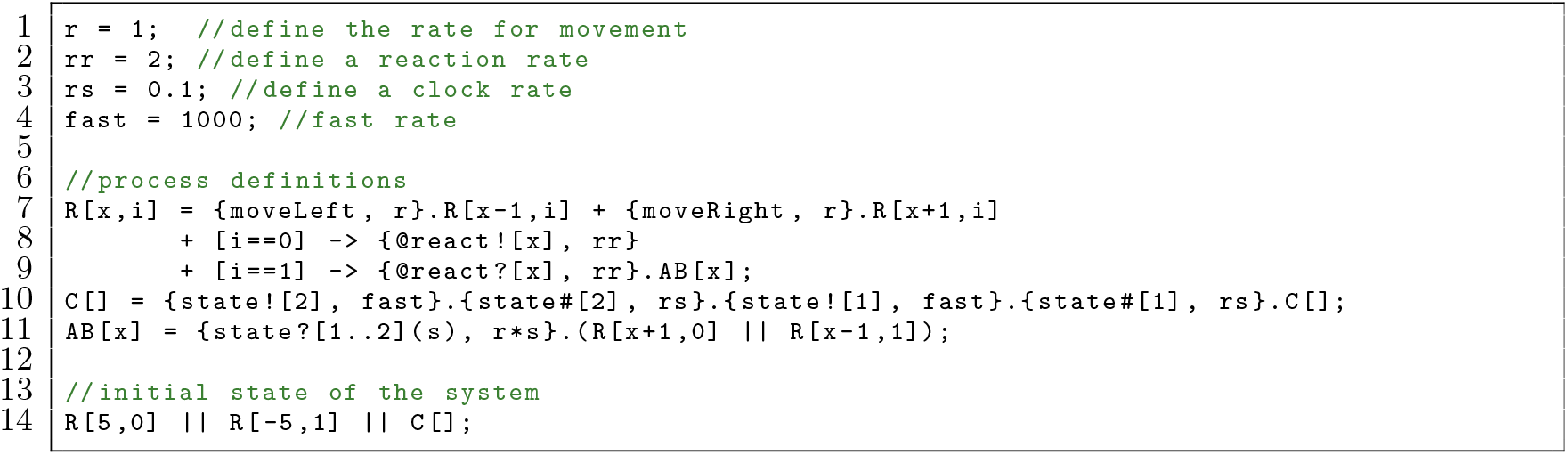

Thus far, these examples have used strings as handshake or beacon channel names which transmitted a single parameter value. These names can also be comma-separated lists, where each entry is an expression of parameters and/or global variables. This allows a process to dynamically change the channel name, and therefore the other processes it can interact with. Likewise, rather than transmitting a single value and receiving a set of values, handshakes and beacons can transmit a comma-separated list of values and receive a comma-separated list of sets. To illustrate with a two-process model where the only possible action is a handshake:

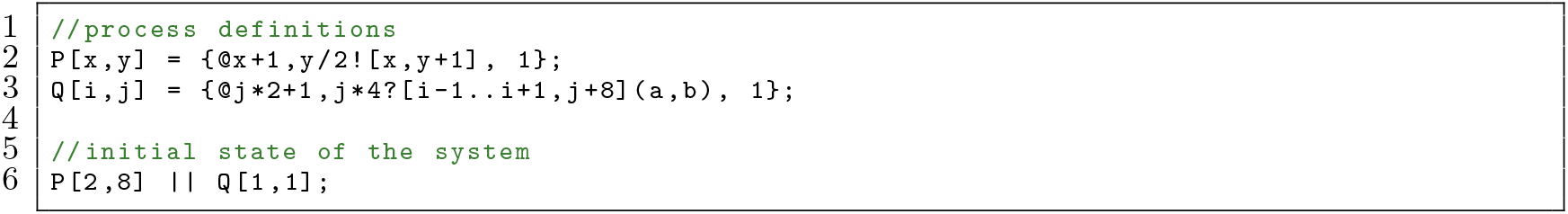

Processes `P` and `Q` will handshake over the channel name `3,4` because `P` transmits the values `2,9` such that `2` lies within the range `i−1‥i+1` (where `i=1`) and `9` is equal to `j+8` (where `j=1`). Process `Q` will then bind the value `2` to variable a and the value `9` to variable `b`. However, if the handshake send by process `P` were changed to {@x+2,y/2![x,y+1], 1} or {@x+1![x,y+1], 1}, the handshake no longer takes place as the channel names do not match. Likewise, {@x+1,y/2![x,y+1,x], 1} would also not result in a handshake as the comma-separated list of parameters must be of the same length between the handshake send and handshake receive.

The ability to use comma-separated lists of values and expressions for handshakes and beacons is particularly important for models where multiple dimensions are considered. The following example returns to the bimolecular reaction *A* + *B* ↔ *AB*:

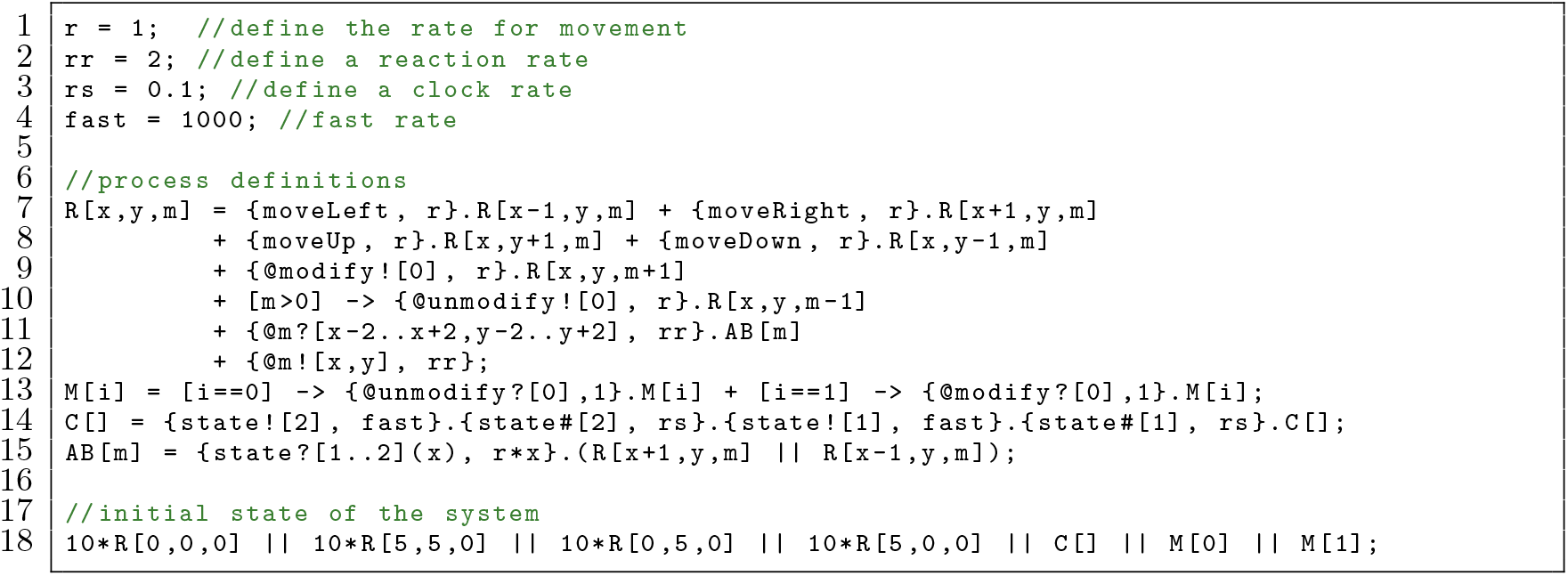

In this model, a reactant `R` has x- and y-coordinates defined by its parameters `x` and `y`, as well as a number of times it was modified `m`. There is a process `M` with parameter `i` that will remove a modification from `R` if `i=0` and add a modification to `R` if `i=1` (Line 13). A reactant `R` can diffuse (Lines 7-8), and it can be modified or unmodified via a handshake with `M` which increments or decrements the value of its parameter `m` (Lines 9-10). The value of `m` is used as a channel name to transmit the x- and y-values of `R` so that only reactants that are nearby and have the same number of modifications can react to create `AB` (Lines 11-12).

The models in this subsection were necessarily arteficial to introduce the Beacon Calculus by way of simple examples, but the following three subsections show the Beacon Calculus applied to three different areas of biological research: DNA replication, DNA damage response, and multisite phosphorylation ultrasensitivity. These diverse examples demonstrate the breadth of applications for the Beacon Calculus and each example showcases a key feature. In the DNA replication model, replication forks use beacons to efficiently coordinate which parts of the chromosome have and have not been replicated. The DNA damage model uses parameters to keep count of damage and repair proteins, showing how to model a population of cells that grows and changes over time. The multisite phosphorylation model shows how receiving a set of possible values in a handshake receive can reduce the number of process definitions required in a model.

### DNA Replication

The mechanisms underlying DNA replication are detailed in a recent review [27] and are briefly summarised here to provide the necessary background for the model. In budding yeast (*S. cerevisiae*), DNA replication initiates during S-phase of the cell cycle from discrete sites on the chromosome known as origins of replication. To maintain genomic integrity, the genome must be fully replicated exactly once per cell cycle. The regulatory mechanism responsible for maintaining this integrity uses an origin recognition complex that binds to the origin and recruits additional proteins to form a pre-replicative complex (pre-RC) in late M-phase and G1-phase when cyclin-dependent kinase (CDK) levels in the cell are low. By the the end of G1, CDK levels have risen (and remain high for the remainder of the cell cycle) so that no new origins can assemble a pre-RC. Those origins that have assembled a pre-RC by S-phase are said to be licensed. The chromosome is replicated when these licensed origins “fire” during S-phase to create bidirectional replication forks that travel along the chromosome in opposite directions from the origin. Forks terminate when they meet a fork travelling in the opposite direction or reach the end of a chromosome.

A random subset (but typically not all) of a chromosome’s origins initiate replication in S-phase and multiple forks can be active at the same time (Fig. 1a). The probability with which origins are licensed is not uniform; some origins are more likely to assemble a pre-RC than others. In addition, of those origins that are licensed, some fire characteristically early in S-phase while others tend to fire late. Therefore, DNA replication is a stochastic process: the set of active origins and the origins responsible for replicating each position on the chromosome will differ from cell-to-cell. Despite this heterogeneity, DNA replication is a remarkably reliable process where errors such as replication fork collapse are rare.

**Figure 1:**
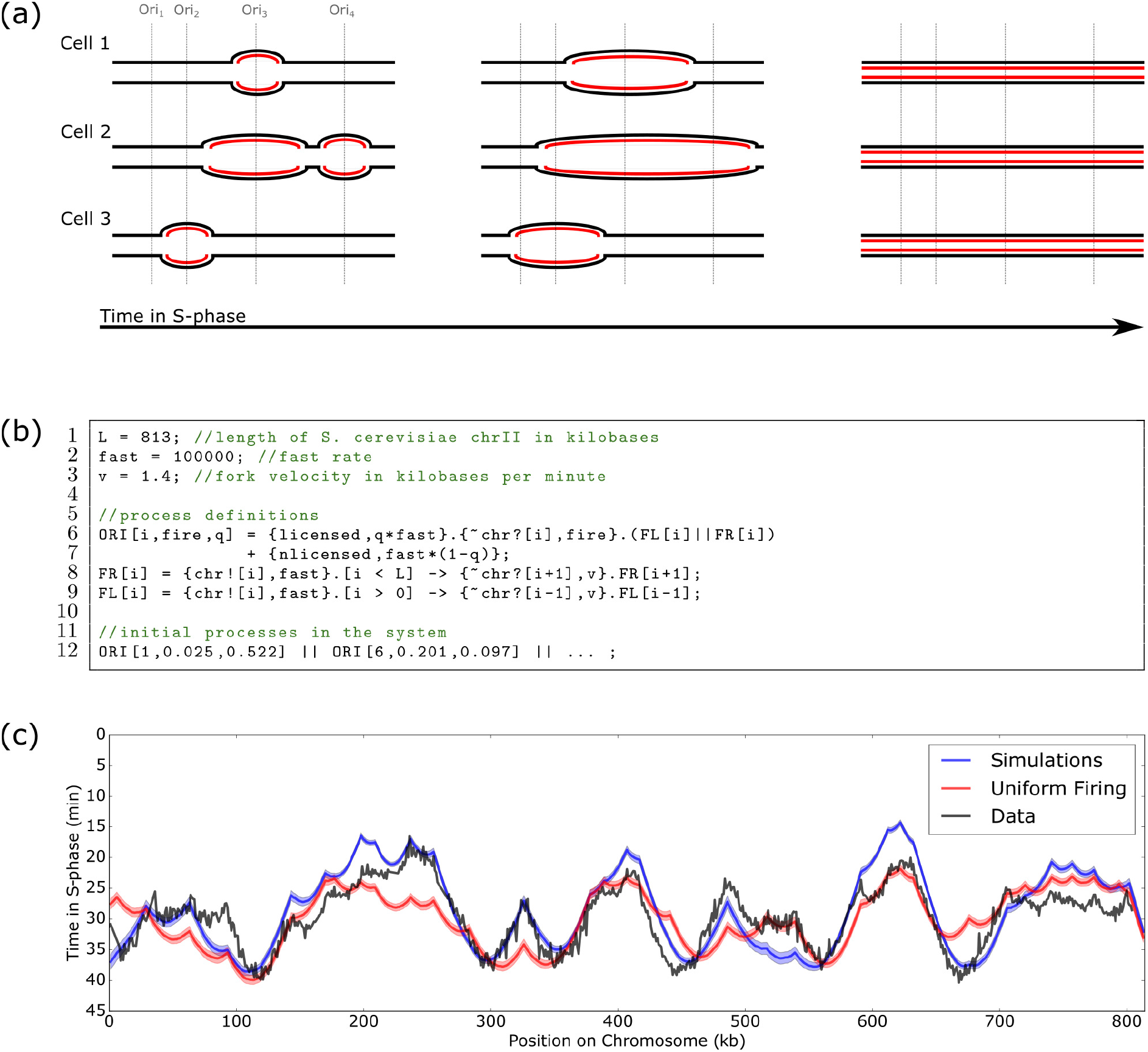
Replication timing from Beacon Calculus simulations. **(a)** Diagram of ongoing DNA replication in the same chromosome segment of three different cells. Replication can begin from four discrete locations (origins of replication). Each cell successfully replicates its DNA despite having different patterns of origin activation. **(b)** DNA replication model written in the Beacon Calculus. **(c)** Curves showing the mean time at which each position on *S. cerevisiae* chromosome II was replicated, taken from the Beacon Calculus model where each origin has a licensing probability and firing time taken from [28] (blue), the Beacon Calculus model where all origins are licensed and have the same firing rate of 0.015 (red), and experimental data from [29] (grey). The Beacon Calculus results are averaged over 500 simulations and shaded regions show the standard error of the mean. The system line has been truncated for clarity (see Section S3).

The stochastic nature of DNA replication makes it well-suited to modelling with the Beacon Calculus: the difference in behaviour between simulations mirrors the heterogeneity between replicating cells, and communication via beacons enables origins and forks to keep track of which chromosomal positions have been replicated. DNA replication is simulated using the Beacon Calculus model in Fig. 1b. The model is comprised of three process definitions: rightward-moving forks `FR`, leftward-moving forks `FL`, and origins of replication `ORI`. The chromosome is of length `L`, and each of these three processes have a single parameter `i` which is taken to be a position on the chromosome between 1 and `L`. Origins have two additional parameters: a licensing probability `q` and a firing rate `fire`. The processes keep track of which chromosomal positions have already been replicated by using beacons: When a fork replicates position `i`, it launches a beacon on channel `chr` with parameter `i`.

The behaviour of an `ORI` process is encoded on Line 6 of Fig. 1b. An origin is licensed or not licensed, which is modelled by the choice between the actions `licensed` and `nlicensed`. If the origin is not licensed, the origin can perform no further actions; it is said to be deadlocked. If the origin is licensed, it fires by performing a beacon check action on channel `chr` at its position `i` to ensure that it only fires if that chromosomal position has not yet been replicated by another fork. Once the origin fires, the `ORI` process continues on as two parallel processes: a rightward-moving fork (`FR`, Line 8) and a leftward-moving fork (`FL`, Line 9). The forks first launch a beacon on channel `chr` with their position to indicate to all other forks and origins that the position has been replicated. After launching the beacon, forks use a gate to ensure they have not yet reached the end of the chromosome. If they have not, the forks verify that the position ahead has not yet been replicated by performing a beacon check on that position. If there is no active beacon at that location, the position has not yet been replicated and the fork moves forward by increasing (for `FR`) or decreasing (for `FL`) the position parameter `i`. Like all processes in the Beacon Calculus, fork movement is stochastic, but forks will tend to the same average velocity over long timescales. Replication has finished when all processes have deadlocked. The initial processes in the system (Line 12) are all `ORI` processes with positions corresponding to 34 known origin locations on *S. cerevisiae* chromosome II [30]. As shown in Fig. 1c, when the initial processes in the system are set to be origins with the positions, licensing probabilities, and firing rates from the literature [28], simulations of the Beacon Calculus model in Fig. 1b give good agreement (*R*^2^ = 0.76) with established replication timing profiles [29].

The simplicity of the DNA replication model in the Beacon Calculus makes it quick and simple to test biological hypotheses. For example, the licensing probability, affinity for firing factors, and the spatial distribution of origins across the chromosome will all have an effect on the replication timing profile; the model can be easily modified to investigate the effect of the spatial distribution of origins alone. The red curve in Fig. 1c shows the timing profile for a modified version of the model where all origins are licensed and the firing rate of all origins is set to the same value. While the timing profile does not match the data as well (*R*^2^ = 0.49), the main features of the replication profile are still captured. This suggests that the primary factor influencing the replication timing profile is the spatial distribution of origins, and that an origin’s affinity for licensing and firing factors play a more minor role. As shown in Fig. 4, making other minor modifications to the the Beacon Calculus model in Fig. 1b allows for modelling cooperative origin firing and replication fork progression through fork barriers. More broadly, this modelling strategy is applicable to coordinated movement by biological components within a reference frame.

### Cellular Response to DNA Damage

To show how the Beacon Calculus can be used to model systems at the population level, this section models the *E. coli* DNA methylation damage system studied in [31]. The effective identification and repair of DNA damage is essential to genome integrity. Unrepaired methylation damage is particularly cytotoxic and mutagenic [32]. In *E. coli*, DNA methylation damage is repaired by the Ada methyltransferase protein: Ada repairs the damage by transferring a methyl group from *O*^6^-Methylguanine or *O*^4^-Methylthymidine to itself [33]. The resulting methylated Ada (meAda) significantly upregulates transcription of the *ada* gene, creating a positive feedback loop that increases Ada levels. This leads to a spike in Ada level following DNA repair which is reduced back to basal levels over generations by successive cell divisions (Fig. 2a).

**Figure 2:**
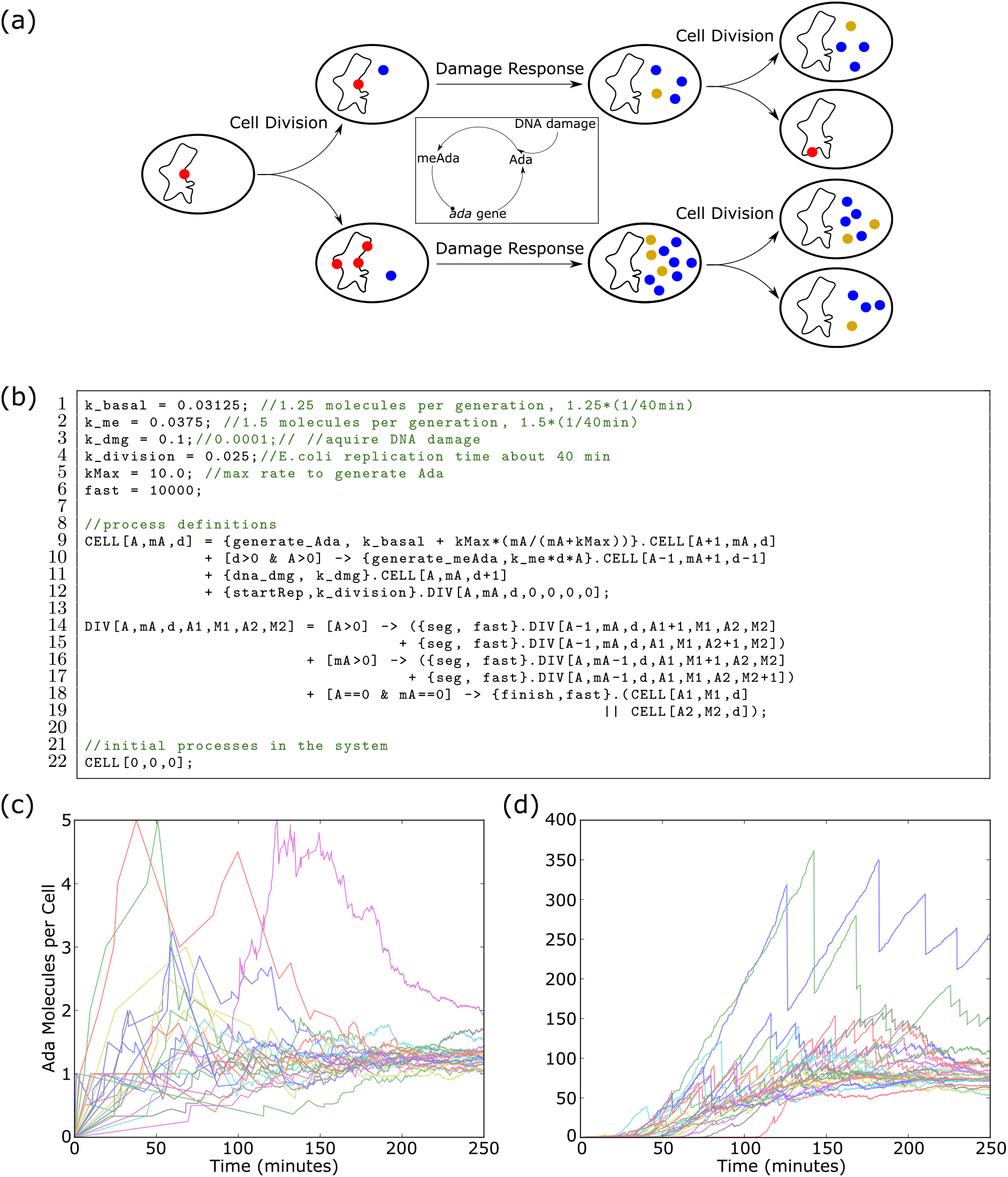
DNA damage from Beacon Calculus simulations. **(a)** Cells undergo DNA damage (red) and may carry it forward for generations before an Ada molecule (blue) is generated to repair it. Ada is methylated (gold) as it repairs DNA damage creating a positive feedback loop whereby methylated Ada upregulates transcription of the *ada* gene. Ada levels are reduced through successive cell divisions. **(b)** DNA damage model written in the Beacon Calculus. **(c-d)** Average total Ada and meAda per cell over time. Each trace corresponds to a simulation of a growing population of cells for **(c)** low DNA damage (*k*_*dmg*_ = 0.0001) and **(d)** high DNA damage (*k*_*dmg*_ = 0.01). In each of panels **(c-d)**, 25 simulations are shown. Values for `k_basal` and `k_division` are from [31] while `k_me` is from [35]. Values for `k_dmg` and `kMax` were approximated based on the results in [31].

A cell keeps Ada levels low in order to perform a delicate balancing act: Excessive Ada levels are thought to be cytotoxic [34], but the cell must still produce enough Ada to repair DNA methylation damage in a timely fashion. This is accomplished by expressing the *ada* gene at very low levels such that on average only one Ada protein is produced per generation [31]. Such a low rate of production means that due to stochasticity, DNA damage may go unrepaired for one or more generations before an Ada protein is produced to repair it (Fig. 2a).

A stochastic model can provide insight into this repair system by showing the Ada response in rare but important situations where DNA methylation damage has gone unrepaired for several generations (see, for example, the complementary model in [35]). By varying the DNA damage rate, a model can also predict how the repair system responds to both high and low rates of DNA methylation damage. The Beacon Calculus makes modelling this system straightforward by representing an *E. coli* cell as a process that can repair DNA damage and divide into two daughter cells (Fig. 2b). The cell process keeps track of DNA damage and Ada levels using parameters and the value of these parameters can scale the rate at which the process performs certain actions.

The process `CELL` is defined on Line 9 of the Beacon Calculus model in Fig. 2b. `CELL` has parameters that keep track of three quantities: the number of Ada molecules (`A`), the number of methylated Ada molecules (`mA`), and the number of sites where DNA has been damaged (`d`). The cell can generate an Ada molecule with action `generate_Ada` (Line 9). The parameter `mA` is used in the rate calculation of this action so that Ada is generated at a basal rate if `mA=0`, but the rate scales to saturation with the value of `mA` to reflect the upgregulation of the *ada* gene by meAda. If the cell has DNA damage and at least one Ada molecule to repair it, `CELL` can fix the damage by first performing action `generate_meAda` and then converting Ada to meAda (Line 10). Damage repair requires interaction between Ada and a methylated base, so the rate of this action scales with the value of `d` and `A`. The cell’s DNA is damaged at a static rate (Line 11) which increments parameter `d`.

The cell can divide at the mean rate of replication for *E. coli* cells (Line 12). When cell division begins, the `CELL` process carries on as a new process `DIV` for a dividing cell. The `DIV` process (Line 14) encodes how Ada, methylated Ada, and damage are segregated between two daughter cells. In addition to the original three parameters of the dividing cell, this process has four additional parameters: the amount of Ada and meAda that segregates to one daughter cell (`A1` and `M1`, respectively) and the amount of Ada and meAda that segregates to the second daughter cell (`A2` and `M2`). These new parameters each start at zero (Line 12). For each Ada and methylated Ada molecule in the parent cell (Lines 14 and 16, respectively) a random choice is made as to which daughter cell inherits the protein. When a choice has been made for each molecule of Ada and meAda, the `DIV` process starts two new daughter `CELL` processes (Lines 18-19).

The initial condition for each simulation is a single cell with no Ada and no DNA methylation damage (Line 22). As the cell divides, the system is comprised of an exponentially growing population of the initial cell’s descendents. Computing the average Ada per cell for this exponentially growing colony shows that the amount of Ada stays near the basal average amount of 1.25 molecules per cell when the rate of DNA damage is low (Fig. 2c, highest spike at 5 Ada molecules per cell). Some colonies exhibited sharp spikes in Ada levels caused by DNA damage that had gone unrepaired for several generations. However, this happened infrequently and the elevated Ada levels tended back towards zero as the Ada was diluted by successive cell divisions. When the rate of DNA damage was high, the spikes in Ada level were higher in magnitude (Fig. 2d, highest spike at 350 Ada molecules per cell). In addition, Ada levels stayed elevated over time and did not tend back towards zero. These observations are qualitatively consistent with the results from [31] of Ada levels in individual *E. coli* cells under both high and low DNA damage conditions.

Communication between processes in the Beacon Calculus means that the model can be easily extended to incorporate cell-to-cell interactions or cell-to-environment interactions using handshakes and beacons. More generally, the Beacon Calculus makes it simple to model a growing and changing population. While this example focused on how a population of cells responds to DNA damage, a similar approach can be taken to model more diverse applications such as the spread of disease through a population.

### Multisite Phosphorylation

Cellular signalling relies on post-translational modifications and, in many instances, substrates are modified on multiple sites. This is thought to confer specific information processing functions such as switch-like responses [36–38] (see [39] for a review). One example is the reversible phosphorylation of membrane-anchored receptors or adaptors by extrinsic kinases and phosphatases, which applies to a large class of receptors known as non-catalytic tyrosine-phosphorylated receptors (NTRs) of which the T-cell antigen receptor (TCR) is a member [40]. NTRs are known to have multiple phosphorylation sites (20 in the case of TCRs) and are phosphorylated and dephosphorylated by kinases and phosphatases that are also confined to the plasma membrane. Given that these receptors often control cellular responses, their phosphorylation is tightly regulated and consequently, will be highly sensitive to the relative concentration or activity of their regulating kinases and phosphatases. This leads to so-called ultrasensitivity, where an input signal produces very little output signal as long as the input remains below a certain threshold, but causes a high output signal once the threshold is exceeded. This results in a sigmoidal input-output curve, typically with a very steep inflection. Ultrasensitivity represents an important way in which biomolecular processes remain robust to noise.

The Beacon Calculus model in Fig. 3 is similar to, and inspired by, the model by Dushek *et al.* who modelled the phosphorylation of TCRs when they were phosphorylated by kinase Lck and dephosphorylated by phosphatase CD45 [41]. The model shown here corroborates the authors’ findings: In order to achieve ultrasensitivity, an enzyme must dwell for a short period after modifying the phosphorylation of a receptor.

**Figure 3:**
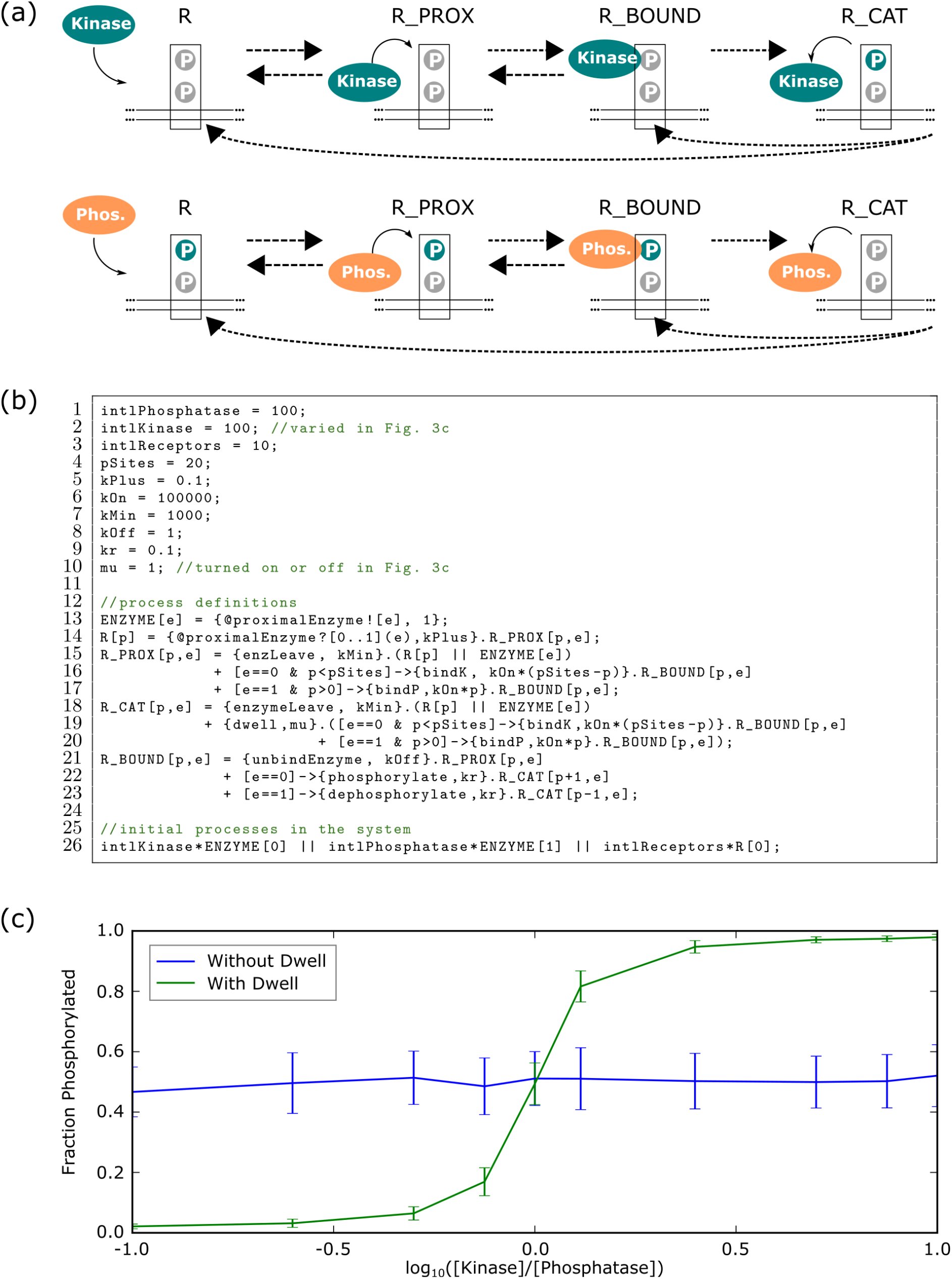
T cell receptor ultrasensitivity from Beacon Calculus simulations. **(a)** An enzyme enters the proximity of a receptor, binds to the receptor, and phosphorylates the receptor if the enzyme is a kinase or dephosphorylates the receptor if the enzyme is a phosphatase. **(b)** T cell receptor model written in the Beacon Calculus with parameters taken from [41]. **(c)** The fraction of phosphorylated receptors is ultrasensitive to the relative concentration of kinase and phosphatase when the enzyme dwells after modifying a receptor (green) but loses ultrasensitivity if the dwell is removed (blue). Points (shown with standard errors) are the average of 50 simulations taken after the system reaches a steady state.

Each TCR can be phosphorylated 20 times. When an enzyme enters the proximity of a receptor, it can either bind to the receptor or leave (Fig. 3a). The enzyme can phosphorylate the receptor if it is a kinase or dephosphorylate the receptor if it is a phosphatase. Once the enzyme phosphorylates or dephosphorylates the receptor, there is a period of inactivity (or a dwell) before the enzyme can bind to the receptor again. The number of phosphorylation sites, together with the action of two types of enzyme, leads to a high number of distinct species in the system; this can make a differential equation model cumbersome to write down and integrate. The Beacon Calculus makes modelling this system straightforward by representing enzymes and receptors as processes, whereby receptor processes keep track of their phosphorylation and the type of enzyme bound to them using parameters.

A model in the Beacon Calculus for TCR phosphorylation is shown in Fig. 3b. Each `ENZYME` process (Line 13) has parameter `e`, whereby the enzyme is a phosphatase if `e=1` or else it is a kinase if `e=0`. A receptor process `R` (Line 14) has parameter `p` which keeps track of the number of times the receptor has been phosphorylated. An enzyme enters the proximity of a receptor via a handshake on channel `proximalEnzyme`, whereby the enzyme transmits its parameter e to the receptor to indicate whether it is a kinase or a phosphatase. After the handshake, the enzyme deadlocks while the receptor carries on as a new process `R_PROX` (Line 15) that encodes the behaviour of a receptor with an enzyme in close proximity. The reverse reaction can occur if `R_PROX` performs action `enzLeave` (Line 15) where `R_PROX` then carries on as `R` and `ENZYME` in parallel. If the enzyme is a kinase and the receptor is not fully phosphorylated, the enzyme can bind at a rate proportional to how many sites on the receptor are unphosphorylated (Line 16). If the enzyme is a phosphatase, the enzyme binds at a rate proportional to how many sites on the receptor are phosphorylated (Line 17). When `R_PROX` binds an enzyme, it carries on as process `R_BOUND`. In this new process, the enzyme can either unbind (Line 21), phosphorylate the receptor if the bound enzyme is a kinase (Line 22), or dephosphorylate the receptor if the bound enzyme is a phosphatase (Line 23). If the enzyme phosphorylates or dephosphorylates the receptor, the bound receptor `R_BOUND` carries on as process `R_CAT` in which the enzyme is proximal to the receptor but briefly inert. The enzyme can either leave (Line 18) or rebind once the inert period is over (Line 19-20).

The above model is similar to that of [41], and the results agree with the authors’ findings (Fig. 3c). When the enzyme dwells after modifying the phosphorylation of a receptor, the fraction of receptors that are phosphorylated is ultrasensitive with respect to the relative concentration of kinase and phosphatase; it displays switch-like behaviour. If the dwell is removed and all other parameter values are kept constant, the ultrasensitivity is lost. While the Beacon Calculus is able to reproduce an established model in only a few lines of code, the language also makes it simple to expand upon the model. For example, the model in Fig. 3b can be extended to model groups of receptors on different areas of the membrane. A group of receptors can use beacons to signal a state change in that group which can cause other groups located elsewhere to respond.

### Extensions to the DNA Replication Model

To demonstrate the flexibility of models written in the Beacon Calculus, the DNA replication model from Fig. 1 is extended to include two topics of interest from the field: cooperative origin firing and the effect of a replication fork barrier.

It has been hypothesised that the probability of a replication origin firing increases if a nearby origin fires [42]. This may be due to stoichiometrically limiting firing factors which are more likely to interact with an origin if they have already interacted with another origin nearby. The Beacon Calculus model in Fig. 4a extends the DNA replication model so that when an origin fires, it launches a beacon on channel `coop` transmitting its location to induce firing of nearby origins. If an origin has not yet fired, it can fire at rate `fire` which is taken to be the origin’s base affinity for firing factors (Line 7). The modified model includes an additional pathway to firing where an origin can receive a beacon on channel `coop` that is transmitting a parameter within 50 kb of the origin’s location (Line 8). The rate at which this beacon is received is inversely proportional to the distance between the origin and the transmitted parameter. If the beacon is received, the origin fires, launches its own beacon on channel `coop` transmitting its position to other origins, and starts two replication forks from its position. Therefore, origin firing is either due to the origin’s natural affinity for firing factors or due to another origin firing nearby (if there is one).

**Figure 4:**
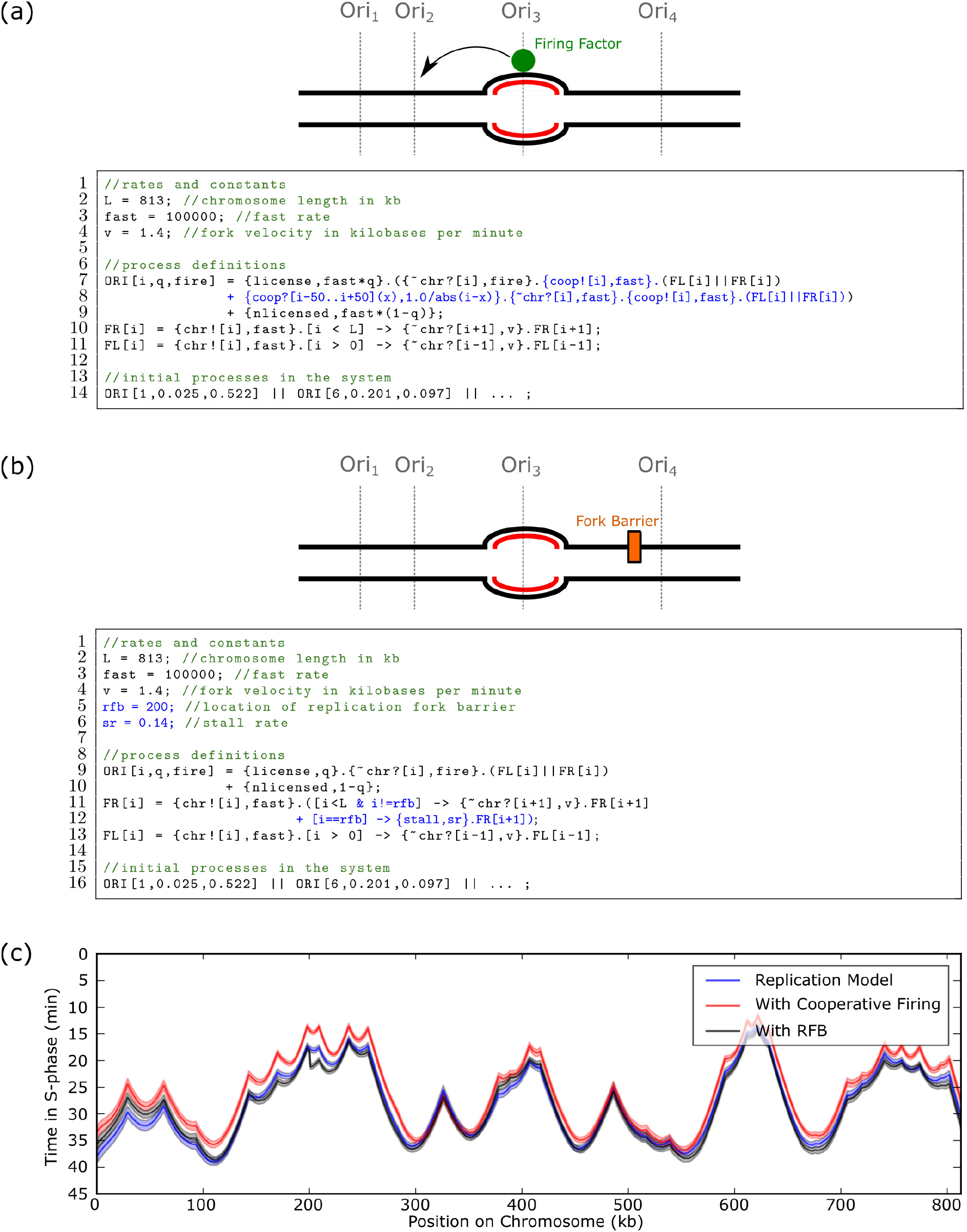
Extensions to the DNA Replication Model. The DNA replication model in Fig. 1b is modified to include either **(a)** cooperative origin firing or **(b)** fork progression through a replication fork barrier. All changes to the model in Fig. 1b are highlighted in blue. **(c)** Results from Fig. 1b compared with the simulated models from **(a)** and **(b)** where each curve is the average over 500 simulations. Shading shows the standard error of the mean. The system line has been truncated for clarity (see Section S3). The stall rate `sr` was chosen to be on average ten times slower than it takes a fork to move 1 kb.

When proteins bind tightly to DNA, they may act as a replication fork barrier (RFB) that can stall replication forks moving in a particular direction. One role of RFBs is preventing collisions between replication and transcription machinery [43]. This is incorporated into the Beacon Calculus model as shown in 5b. First, the location of the fork barrier is specified on the chromosome (Line 5) along with the rate of the stall (Line 6). If a rightward moving fork makes it to this position, it stalls (Line 12) before it ultimately recovers and continues stepping.

Simulations of the models shown in Fig. 4a-4b are shown in Fig. 4c. The replication fork barrier causes a sharp change to the timing profile near the location of the replication fork barrier at chromosomal coordinate `i=200` while the cooperative firing behaviour makes the whole chromosome replicate slightly earlier. However, the additional parameters added in these two models were not fit to data; these two extensions are only intended to demonstrate the ease with which Beacon Calculus models can be extended. With further parametrisation, however, these extensions can be useful in making meaningful biological predictions about DNA replication systems.

## Discussion

Process calculi are a natural framework in which to model biological systems, but they are an underutilised tool within systems biology; to the authors’ knowledge, process calculi have never before been applied to DNA replication, DNA methylation damage, or receptor ultrasensitivity. The Beacon Calculus makes it quick and easy to create models of systems where processes can change both their actions and interactions over time. Beacons make it simple for a process to influence the actions of all other processes in the system. This paper has shown how this paradigm is used to model both the complex behaviour of cells and macromolecular structures in only a few lines of code.

A language that makes it simple and concise to encode biological models has advantages beyond saving time: it changes the way the tool is used. Simplicity increases confidence that the user has actually encoded what they think they have encoded and have not introduced bugs into the model. It also leads to models that are easy to change, modify, and extend. This flexibility encourages experimentation where hypotheses can be rapidly tested, and any conclusions drawn from laboratory experiments investigated, to ensure that they are consistent with the biological data.

The features in the Beacon Calculus are all geared toward models that are quick to encode and easy to modify. As shown above, the DNA replication model in Fig. 1b can be modified to include features of interest from the literature such as cooperative origin firing and a replication fork barrier that stalls replication forks at a particular chromosomal coordinate. The flexibility of the Beacon Calculus means that these changes are straightforward to incorporate and come at the expense of only one or two lines of code.

While this paper has shown that the Beacon Calculus can easily produce flexible and concise models of biological systems from the current literature, it is not appropriate for every task. Section S2 compares the Beacon Calculus with the stochastic *π*-calculus [15, 16, 18], Kappa [47–49], Bio-PEPA [9], BioNetGen [44, 45], PySB [46], ML-Rules [50, 51], and Simmune [52, 53]. For each of these tools, examples are described where they may be more appropriate than the Beaocn Calculus. In general, rule-based languages may be the better choice for applications where the complex, combinatorial assembly of biomolecules is important. This is particularly important for applications involving large protein-protein interaction networks and modification of species by ligands. In addition, while it is possible to create species within a compartment with Beacon Calculus parameters, tools such as Bio-PEPA, ML-Rules, and Simmune deal with this much more naturally. The Beacon Calculus finds its niche in applications where system components must be able to easily coordinate with each other or with a global reference frame (such as in the DNA replication model) or adapt behaviour in response to complex and changing environmental conditions (such as a cell responding to DNA damage or multisite phosphorylation).

There are many applications throughout biology where the Beacon Calculus can be an ideal tool for modelling and simulation. This paper illustrated three examples from cell biology and molecular biology, but modelling at the population level is possible as well. A stochastic version of the SIR model for a population’s response to an infectious disease would be straightforward: each individual is a process, whether they are susceptible, infected, or recovered from an infection is kept track of with a parameter, a response to nearby individuals could be modelled using the ability of handshake receives to accept a range of parameters, and beacons could be used to signal some state change within a city or area as the disease evolves. There are a wide range of applications within biology, and while the Beacon Calculus was developed for biological applications, there is nothing biology-specific in the language; it can be used for applications in engineering and other fields.

One of the biggest challenges in creating a simulation tool is ensuring the user is simulating what they think they are simulating; if the user has made an error encoding the model, this can lead to incorrect conclusions being drawn about the underlying biology. An advantage of process algebras is that the language’s semantics, together with automated theorem proving techniques, can be used to prove whether a certain combination of actions is ever possible in the model. In the DNA replication model, for example, a user may wish to verify that replication forks cannot step through each other in the model that they have encoded. If this action is possible, then there is an error in the model and the simulation results will not accurately reflect the biological reality. A planned extension of the bcs tool is allowing the user to specify certain actions or properties that should not be allowed in the model. The tool will check these properties before beginning the simulations to ensure that they are not possible, giving the user greater confidence in the validity of the result.

The Beacon Calculus is a language that makes it fast and easy to encode concise, flexible models of biological systems. It is particularly well-suited for systems where interactions between components change over time, where components need to change the state of many other components, or where components need to respond to events happening within a certain region. Its breadth is demonstrated by creating models of DNA replication and DNA damage repair from the literature, as well as creating a stochastic version of an established deterministic multisite phosphorylation model. To support the language, a contribution of this work is an open-source simulator called bcs which, together with the provided examples, makes it easy for users to create and simulate their own models.

## Materials and methods

An open-source Beacon Calculus simulator (bcs) is provided to simulate models written in the Beacon Calculus (https://github.com/MBoemo/bcs.git). The software uses a modified Gillespie algorithm to simulate paths through the model [26]. For each simulation, the software outputs a table of actions sorted in order of ascending time. Each row specifies a time, the action performed at that time, and the process that performed the action (as well as its parameter values at the time when the action was performed). While there is basic plotting capability included with the software, the output was designed to be easy to parse so that it can be reformatted into plots that are appropriate for the biological system being modelled. For the results in this paper, the Beacon Calculus output has been reformatted into plots that are common in the examples’ respective fields. To make it clear how to use the bcs software to simulate biological models, all of the examples in this paper are written in bcs source code. Benchmarks for the run time of simulations are specified in Section S4.

## Acknowledgments

The authors are grateful to Omer Dushek (Sir William Dunn School of Pathology, University of Oxford) and Stephan Uphoff (Department of Biochemistry, University of Oxford) for helpful discussions and guidance on the multisite phosphorylation and DNA damage examples, respectively. This work was supported by Biotechnology and Biological Sciences Research Council (https://bbsrc.ukri.org/) grant BB/N016858/1 and Wellcome Trust (https://wellcome.ac.uk/) Investigator Award 110064/Z/15/Z to CAN. Additional funding and support is provided by the St. Cross College Emanoel Lee Junior Research Fellowship to MAB, as well as funds to MAB from the Department of Pathology, University of Cambridge.

## Supporting information

**S1 Appendix. Language Definition.** A formal definition of the Beacon Calculus language, including the grammar and structural operational semantics.

**S2 Appendix. Comparison with Other Methods.** Case studies and discussion comparing the Beacon Calculus to three other established methods: the stochastic *π*-calculus, Kappa, PEPA, BioNetGen, PySB, ML-Rules, and Simmune.

**S3 Appendix. DNA Replication Model.** Further details and discussion relating to the DNA replication model.

**S4 Appendix. Benchmarks.** The average computation time needed to compute the DNA replication model, the DNA damage model, and the multisite phosphorylation model.

**S5 Appendix. Additional Example: Kinesin Stepping Down a Microtubule.** An additional introductory example that describes the movement of kinesin down a microtubule. This example complements the example in Language Overview.

## Software and Data Availability

The Beacon Calculus simulator is available at https://github.com/MBoemo/bcs.git.

## Notes

https://github.com/MBoemo/bcs.git

