## Supplemental Information for "The Beacon Calculus: A formal method for the flexible and concise modelling of biological systems"

### Supplementary Information

Michael A. Boemo[1*] Luca Cardelli[2] Conrad A. Nieduszynski[3]

[1]Department of Pathology, University of Cambridge
[2]Department of Computer Science, University of Oxford
[3]Genome Damage and Stability Centre, University of Sussex

#### S1 Language Definition

This section complements the informal description given in the Language Overview subsection of the main text by formally specifying the grammar (Section S1.1 and structural operational semantics (Section S1.2) of the Beacon Calculus; this dictates both how to write models in the Beacon Calculus and how they are evaluated.

##### S1.1 Grammar

Beacon Calculus models consist of variable definitions, followed by process definitions, followed by a system line that specifies the initial state of the system. The context-free grammar of a Beacon Calculus model is specified below. For clarity, terminals in the grammar are coloured in red while nonterminals are enclosed in angled brackets.

$$
\begin{aligned}
\langle Model \rangle ::= & [\langle VariableDef_1 \rangle;\ \langle VariableDef_2 \rangle;\ ...\ ;\ \langle VariableDef_N \rangle;] \\
& \langle ProcessDef_1 \rangle;\ \langle ProcessDef_2 \rangle;\ ...\ ;\ \langle ProcessDef_N \rangle; \\
& \langle SystemLine \rangle;
\end{aligned}
$$

$$
\begin{aligned}
\langle VariableDef \rangle ::= & \langle Id \rangle = \langle Int \rangle \\
& | \langle Id \rangle = \langle Float \rangle
\end{aligned}
$$

$$
\langle ProcessDef \rangle ::= \langle Id \rangle [\langle IdList \rangle \mid \epsilon] = \langle Process \rangle
$$

$$
\begin{aligned}
\langle Process\rangle ::= {} & \epsilon \\
& |\langle Id\rangle[\langle ExpressionList\rangle|\epsilon\rangle] \\
& |\langle Process\rangle+\langle Process\rangle \\
& |\langle Process\rangle\|\langle Process\rangle \\
& |\{\langle Action\rangle,\langle Expression\rangle\}.\langle Process\rangle \\
& |[\langle Condition\rangle]->\{\langle Action\rangle,\langle Expression\rangle\}.\langle Process\rangle
\end{aligned}
$$

$$\langle SystemLine\rangle ::=[\langle Int\rangle*]\langle Id\rangle[\langle NumList\rangle|\epsilon][\|\langle SystemLine\rangle]$$

$$\langle NumList\rangle ::= (\langle Int\rangle|\langle Float\rangle)[,\langle NumList\rangle]$$

$$\langle Action\rangle ::= \{\ (\langle Beacon\rangle|\langle Handshake\rangle|\langle Id\rangle)\ ,\langle Expression\rangle\}$$

$$
\begin{aligned}
\langle Handshake\rangle ::= {} & @\langle Channel\rangle?[\langle SetList\rangle] \\
& |@\langle Channel\rangle?[\langle SetList\rangle](\langle IdList\rangle) \\
& |@\langle Channel\rangle![\langle ExpressionList\rangle]
\end{aligned}
$$

$$
\begin{aligned}
\langle Beacon\rangle ::= {} & \langle Channel\rangle?[\langle SetList\rangle] \\
& |\langle Channel\rangle?[\langle SetList\rangle](\langle IdList\rangle) \\
& |\sim\langle Channel\rangle?[\langle SetList\rangle] \\
& |\langle Channel\rangle![\langle ExpressionList\rangle] \\
& |\langle Channel\rangle\#[\langle ExpressionList\rangle]
\end{aligned}
$$

$$
\begin{aligned}
\langle Set\rangle ::= {} & \langle Int\rangle \\
& |\langle IntExpression\rangle \\
& |\ (\langle Int\rangle\ |\ \langle IntExpression\rangle)\ ..\ (\langle Int\rangle\ |\ \langle IntExpression\rangle) \\
& |\langle Set\rangle\backslash\langle Set\rangle \\
& |\langle Set\rangle\ \mathtt{U}\ \langle Set\rangle \\
& |\langle Set\rangle\ \mathtt{I}\ \langle Set\rangle
\end{aligned}
$$

$\langle Expression \rangle ::= \langle Int \rangle$
$\quad | \langle Float \rangle$
$\quad | \langle Expression \rangle * \langle Expression \rangle$
$\quad | \langle Expression \rangle / \langle Expression \rangle$
$\quad | \langle Expression \rangle + \langle Expression \rangle$
$\quad | \langle Expression \rangle - \langle Expression \rangle$
$\quad | \langle Expression \rangle \wedge \langle Expression \rangle$
$\quad | abs(\langle Expression \rangle)$
$\quad | sqrt(\langle Expression \rangle)$
$\quad | max(\langle Expression \rangle, \langle Expression \rangle)$
$\quad | min(\langle Expression \rangle, \langle Expression \rangle)$

$\langle IntExpression \rangle ::= \langle Int \rangle$
$\quad | \langle Expression \rangle * \langle Expression \rangle$
$\quad | \langle Expression \rangle / \langle Expression \rangle$
$\quad | \langle Expression \rangle + \langle Expression \rangle$
$\quad | \langle Expression \rangle - \langle Expression \rangle$
$\quad | \langle Expression \rangle \wedge \langle Expression \rangle$
$\quad | abs(\langle Expression \rangle)$
$\quad | sqrt(\langle Expression \rangle)$
$\quad | max(\langle Expression \rangle, \langle Expression \rangle)$
$\quad | min(\langle Expression \rangle, \langle Expression \rangle)$

$\langle Channel \rangle ::= \big(\langle IntExpression \rangle \;\big|\; \langle Id \rangle\big) \; [, \langle Channel \rangle]$

$\langle ExpressionList \rangle ::= \langle IntExpression \rangle \; [, \langle ExpressionList \rangle]$

$\langle SetList \rangle ::= \langle Set \rangle \; [, \langle SetList \rangle]$

$\langle IdList \rangle ::= \langle Id \rangle \; [, \langle IdList \rangle]$

$\langle Relation \rangle ::= \langle Expression \rangle > \langle Expression \rangle$
$\quad | \langle Expression \rangle >= \langle Expression \rangle$
$\quad | \langle Expression \rangle < \langle Expression \rangle$
$\quad | \langle Expression \rangle <= \langle Expression \rangle$
$\quad | \langle Expression \rangle == \langle Expression \rangle$
$\quad | \langle Expression \rangle != \langle Expression \rangle$

$\langle Condition \rangle ::= \langle Relation \rangle$
$\quad | \langle Condition \rangle \& \langle Condition \rangle$
$\quad | \langle Condition \rangle | \langle Condition \rangle$
$\quad | \sim \langle Condition \rangle$

$\langle Int \rangle ::= (-)?(0...9)+$

$\langle Float \rangle ::= \langle Int \rangle.(0...9)*$

$\langle Id \rangle ::= (A...Z|a...z|\_) + (A...Z|a...z|\_|0...9)*$

#### S1.2 Structural Operational Semantics

A set of defined processes with their initial parameter values specified is referred to as a system. In the Beacon Calculus, each system also includes a single database $D$ which is a set of ordered channel-value pairs. This database provides the means to keep track of which beacons are active, and it is updated every time a process launches or kills a beacon. Processes query the database each time they perform a beacon receive or a beacon check action. When a process performs an action, the system (made up of processes and the database) transitions to a new state. The allowed transitions are specified by the structural operational semantics of the calculus, which is expressed by the inference rules in Equations (8)-(18).

To make database updates precise, each concurrent process in the system is paired with a ledger $L$. This ledger is an ordered record of channel-value pairs $(c, i)$ that specifies how that process has modified the database via beacon actions. As both $c$ and $i$ are comma-separated lists, equality and set inclusion between two lists are taken element-wise: $i_1, i_2, ..., i_n$ is equal to $\hat{i}_1, \hat{i}_2, ..., \hat{i}_m$ if $n = m$ and $i_j = \hat{i}_j$ for $j = 1, ..., n$. Likewise, $i \in \Omega$ where $i = i_1, i_2, ..., i_n$ and $\Omega = \Omega_1, \Omega_2, ..., \Omega_m$ if $n = m$ and $i_j \in \Omega_j$ for all $j = 1, .., n$. A ledger is defined by,

$$\langle L \rangle ::= \epsilon \;\big|\; !(c, i).\langle L \rangle \;\big|\; \#(c, i).\langle L \rangle \tag{1}$$

where $\epsilon$ is empty, the prefix operator $!(c, i).L$ records a beacon launch of channel-value pair $(c, i)$ on the ledger, and $\#(c, i).L$ records the beacon kill on the ledger. The database $D$ is a function that maps a ledger $L$ onto a set of channel-value pairs:

$$D(\epsilon) = \phi \tag{2}$$
$$D(!(c, i).L) = D(L) \cup (c, i) \tag{3}$$
$$D(\#(c, i).L) = D(L) \setminus (c, i) \tag{4}$$

Two ledgers can be merged into one with the operation $L_1 \bowtie L_2$. This results in a set of ledgers where the order of $L_1$ is preserved, the order of $L_2$ is preserved, but they are interleaved differently. This is made precise by first defining the prefix extension to a set of ledgers:

$$a.S = \{a.L : L \in S\} \tag{5}$$

Then the merge between two ledgers is given by,

$$a.L_1 \bowtie b.L_2 = a.(L_1 \bowtie (b.L_2)) \cup b.((a.L_1) \bowtie L_2) \tag{6}$$

where the merger of a non-empty ledger with an empty ledger results in a singleton set:

$$\epsilon \bowtie L = L \bowtie \epsilon = \{L\} \tag{7}$$

By updating a ledger as it works and converting that ledger into a database when the database needs to be queried, processes can interact with a global database as necessary in the rules below.

There is a partition on the set of actions $\mathcal{A}$ given by,

$$\mathcal{A} = W \cup B \cup H$$

where $B$ is the set of beacon actions (launch, receive, check, and kill), $H$ is the set of handshake actions (send and receive), and $W$ is the complement of $B \cup H$ in $\mathcal{A}$. (That is, $W$ is the set of non-messaging, or "wait", actions.) Following the stochastic $\pi$-calculus, there is a built-in action $\tau \in W$ such that $\{\tau, r\}$ is time passing at rate $r$ [22]. While recursion is not included below, the handling of recursion in structural operational semantics is standard amonst process algebras (see, for example, the use of a replication operator in the $\pi$-calculus [19]) so it is omitted here for brevity and clarity.

*Prefix:*
$(a \in W)$

$$\frac{}{L, \{a, r\}.P \xrightarrow{\{a,r\}} L, P} \tag{8}$$

$$\frac{}{L, \{c![i], r\}.P \xrightarrow{\{c![i],r\}} !(c,i).L, P} \tag{9}$$

$$\frac{}{L, \{c\#[i], r\}.P \xrightarrow{\{c\#[i],r\}} \#(c,i).L, P} \tag{10}$$

$$\frac{\exists\ (c,i) \in D(L) \text{ such that } i \in \Omega}{L, \{c?[\Omega], r\}.P \xrightarrow{\{c?[\Omega],r\}} L, P} \tag{11}$$

$$\frac{\exists\ (c,i) \in D(L) \text{ such that } i \in \Omega}{L, \{c?[\Omega](x), r\}.P \xrightarrow{\{c?[\Omega](x),r\}} L, P\big|_{x=i}} \tag{12}$$

$$\frac{(c,i) \notin D(L) \text{ for all } i \in \Omega}{L, \{\sim c?[\Omega], r\}.P \xrightarrow{\{\sim c?[\Omega],r\}} L, P} \tag{13}$$

*Choice:*
$(a \in \mathcal{A})$

$$\frac{L, P \xrightarrow{\{a,r\}} \hat{L}, \hat{P}}{L, P + Q \xrightarrow{\{a,r\}} \hat{L}, \hat{P}} \qquad \frac{L, Q \xrightarrow{\{a,r\}} \hat{L}, \hat{Q}}{L, P + Q \xrightarrow{\{a,r\}} \hat{L}, \hat{Q}} \tag{14}$$

*Parallel Execution:*
$(a \in \mathcal{A})$

$$\frac{L, P \xrightarrow{\{a,r\}} \hat{L}, \hat{P}}{L, P \parallel Q \xrightarrow{\{a,r\}} \hat{L}, \hat{P} \parallel Q} \qquad \frac{L, Q \xrightarrow{\{a,r\}} \hat{L}, \hat{Q}}{L, P \parallel Q \xrightarrow{\{a,r\}} \hat{L}, P \parallel \hat{Q}} \tag{15}$$

*Handshake:*

$$\frac{L_1, P \xrightarrow{\{@c![i],r_1\}} \hat{L_1}, \hat{P};\quad L_2, Q \xrightarrow{\{@c?[\Omega],r_2\}} \hat{L_2}, \hat{Q};\quad i \in \Omega}{L_3, P \parallel Q \xrightarrow{\{\tau, r_1 \cdot r_2\}} \hat{L_3}, \hat{P} \parallel \hat{Q} \text{ where } \hat{L_3} \in \hat{L_1} \bowtie \hat{L_2}} \tag{16}$$

$$\frac{L_1, P \xrightarrow{\{@c![i], r_1\}} \hat{L_1}, \hat{P}; \quad L_2, Q \xrightarrow{\{@c?[\Omega](x), r_2\}} \hat{L_2}, \hat{Q}; \quad i \in \Omega}{L_3, P \parallel Q \xrightarrow{\{\tau, r_1 \cdot r_2\}} \hat{L_3}, \hat{P} \parallel \hat{Q}\big|_{x=i} \text{ where } \hat{L_3} \in \hat{L_1} \bowtie \hat{L_2}} \tag{17}$$

*Gates:*
$(a \in \mathcal{A})$

$$\frac{\mathcal{B} = \texttt{True}; \quad L, P \xrightarrow{\{a,r\}} \hat{L}, \hat{P}}{L, [\mathcal{B}] \rightarrow P \xrightarrow{\{a,r\}} \hat{L}, \hat{P}} \tag{18}$$

In the inference rules above, Eqn. (8) specifies that if a non-messaging action is prefixed to a process, that action can be done with no premises; in this case, there are no conditions required. Note that the ledger $L$ is not updated in this case. Equations (9)-(10) pertain to beacon launch and beacon kill actions, respectively. These actions also require no premises, but performing either of these actions updates the ledger $L$. For the beacon launch in Eqn. (9), the ledger is appended with the channel and value pair specified in the action; for the beacon kill action in Eqn. (10), the ledger is appended with channel and value pair that have been removed from the ledger.

Unlike the first three rules, the beacon receive in Eqn. (11) does have a premise: In order to receive any value in set $\Omega$ on channel $c$, a value $(c, i)$ with $i \in \Omega$ must be in the database. If the database contains no such pair, then the process must wait until it does in order to perform the beacon receive action. Eqn. (12) is similar, but introduces variable binding: If the beacon receive should bind a variable $x$, then $x$ is substituted for the value received in the rest of the process. The beacon check action specified in Eqn. (13) is the opposite of a beacon receive: it can only be done if there is no such $(c, i)$ pair in the database, and if there is such a pair, the action cannot be done until each beacon $(c, i)$ for $i \in \Omega$ is killed.

Eqn. (14) shows that the choice operator is exclusive: when a process performs an action, if there was a choice operation on that action and another action, then the system cannot go back and do the other action. In contrast, Eqn. (15) shows that when two processes are in parallel, the transition of one process does not affect the other process. Handshakes, as specified by Eqn. (16), require two processes acting in parallel: one that can do a handshake send $\{@c![i], r_1\}$, and another that can do a handshake receive $\{@c?[\Omega], r_2\}$ with $i \in \Omega$. Both of these conditions must hold to handshake. Otherwise, a process needs to wait to send its handshake until a suitable handshake receive becomes available (and vice versa). If two processes can handshake, a system can perform a handshake action whereby both the sending and receiving processes transition at the same time at rate $r_1 \cdot r_2$. Eqn. (17) is similar but includes a binding variable.

Eqn. (18) specifies that if an action is gated, then the Boolean condition imposed by the gate on the process's parameters must evaluate to true in order for the process to perform the gated action. If the condition evaluates to false, then the process cannot perform the action. These gated actions, as well as the use of beacons in Eqns. (11)-(13), draw inspiration from Dijkstra's notion of guarded commands [10].

### S2 Comparison with Other Methods

#### S2.1 Comparison with the Stochastic $\pi$-Calculus

The $\pi$-calculus is a universal model of computation that allows channel names to be communicated to other processes, allowing networks to dynamically reconfigure their connectivity [18, 19]. As the $\pi$-calculus and its stochastic extension are Turing complete, no language (including the Beacon Calculus) is more expressive. However, while any biological model can be encoded in the stochastic $\pi$-calculus, it is not necessarily convenient to do so. The niche of the Beacon Calculus is in providing a way to encode complex biological models in a simple syntax, making models easier to read and write.

While the $\pi$-calculus does not explicitly provide for asynchronous communication, the behaviour of beacons could be emulated in the $\pi$-calculus through replicated actions. If a new process is started that offers a handshake as a replicated action, then this new process effectively acts like a beacon. This new "beacon process" can be killed by receiving a handshake on another channel that causes it to deadlock. For the DNA replication model presented in Fig. 1, a new process would be started for each chromosomal position where each of these processes would communicate its position on the chromosome via a channel. A replication fork process could emulate the beacon check behaviour through a race condition between stepping forward at a low rate or communicating with one of the beacon processes at a faster rate. While this is possible, it is cumbersome. In the SPiM implementation of the stochastic $\pi$-calculus, for example, a new channel would have to be explicitly declared for each chromosomal position. In addition, the number of processes in the system would scale linearly with the length and resolution of the chromosome being considered. In the Beacon Calculus, the number of distinct processes scales linearly with the number of active replication origins, which is much lower than the number of positions in the chromosome. The number of active processes in the Beacon Calculus also does not depend on the resolution of the model: the model has the same number of active processes regardless of whether replication is simulated at base-pair or kilobase resolution.

In the Beacon Calculus, beacon receives and handshake receives enable a process to receive a set of values on a channel. This is distinct from the stochastic $\pi$-calculus implementations (such as SPiM [21]) where each value that can be received on a channel must be individually specified. An example where this feature is useful is when a process should investigate a particular region and respond to it in some way. Evidence from the DNA replication literature suggests that if an origin of replication fires, this can impact the chance of a nearby origin firing either positively or negatively. As shown in Fig. 4a, the Beacon Calculus from Fig. 1 can be easily modified to include cooperative origin firing. The ability to receive a set of values on a channel makes this simple and concise to express.

#### S2.2 Comparison with Kappa

Kappa is a rule-based language for the simulation of protein-protein interaction networks, although it can be used for other types of chemical reaction networks as well [3, 7, 8]. The language allows a user to equip different chemical species with a number of binding sites and user-specified rules determine how, and at what rate, these binding sites can interact. The authors provide a simulation tool that uses the Gillespie algorithm on this set of rules to simulate a path through the system's state space.

The Kappa language comes with an additional "mini-language" of intervention directives, which make it possible to add, delete, and modify species. These interventions can be triggered by a time point or an amount of a particular species. In addition, the language supports tokens, or global variables that can be modified upon the execution of certain rules, as well as counters that can perform tasks like counting the number of phosphorylated sites on a species. It may be possible to implement models like those presented in this paper in Kappa using intervention directives and tokens, but these would not be straigthtforward to encode and extend. (Tokens and counters, in particular, are listed in the Kappa manual as experimental features.) For example, both the DNA damage model and the T-cell antigen receptor model presented in this paper use parameters to scale the rate at which an individual process will perform a certain action. However, the parameter range would have to be explicitly encoded as a number of binding sites. While it may be simply inconvenient to encode the 20 possible phosphorylation sites in the T-cell antigen receptor model, this becomes intractable in the DNA damage model where cells must keep track of unbonded amounts of Ada. DNA replication dynamics would also be cumbersome to encode in Kappa and extend, particularly with the cooperative origin firing extension from Fig. 4a. In the Beacon Calculus, dealing with these features is straightforward and natural due to the language's syntax and semantics.

While the Beacon Calculus is more suited to some examples such as those presented in the paper, Kappa may be the better choice for simulating more typical protein-protein interaction networks. An example system from the Kappa manual includes three proteins: A, B, and C. A and B can reversibly bind to one another. Protein A can only phosphorylate the first site on Protein C when Proteins A and B are bound. The second site on Protein C can only be phosphorylated after the first site, and only by Protein A which is not bound to Protein B. Fig. S1 shows this model written in the Beacon Calculus (Fig. S1a) and in Kappa (Fig. S1b). In this system, the features in the Beacon Calculus do not make the system easier to encode, and the reaction rules have been implemented using gates. While Fig. S2 shows that the simulated Beacon Calculus and Kappa models (using their respective simulation tools) give the same answer, Kappa is the more appropriate tool for this type of system: writing out the reaction rules is easier to read and more comprehensible than using parameters and gates in the Beacon Calculus.

(a)

```
//rates
on_rate = 0.001;
off_rate = 0.1;
mod_rate = 1;

//process definitions
A[i] = {@bindA![0],on_rate};
B[a] = [a==0] -> {@bindA?[0],1}.B[1]
     + [a==1] -> ({unbindA,off_rate}.(A[0] || B[0]) + {@bindAB![0],on_rate});

C[x1,x2,p1,p2] = [x1==0 & p1==0] -> {@bindAB?[0],1}.C[2,x2,p1,p2]
               + [x2==0 & p1==1 & p2==0] -> {@bindA?[0],1}.C[x1,1,p1,p2]
               + [x1==2] -> {unbindB,off_rate}.(C[1,x2,p1,p2] || B[0])
               + [x1==2 & p1==0] -> {phos_x1_AB,mod_rate}.(C[0,x2,1,p2] || B[1])
               + [x1==1 & p1==0] -> {phos_x1_A,mod_rate}.(C[0,x2,1,p2] || A[0])
               + [x2==1 & p2==0] -> {phos_x2,mod_rate}.(C[x1,0,p1,1] || A[0]);

//system
100*A[0] || 100*B[0] || 1000*C[0,0,0,0];
```

(b)

```
/* Signatures */
%agent: A(x,c) // Declaration of agent A
%agent: B(x) // Declaration of B
%agent: C(x1{u p},x2{u p}) // Declaration of C with 2 modifiable sites

/* Rules */
'a.b' A(x[.]),B(x[.]) -> A(x[1]),B(x[1]) @ 'on_rate' //A binds B
'a..b' A(x[1/.]),B(x[1/.]) @ 'off_rate' //AB dissociation

'ab.c' A(x[_],c[.]),C(x1{u}[.]) ->A(x[_],c[2]),C(x1{u}[2])
       @ 'on_rate' //AB binds C
'mod␣x1' C(x1{u}[1]),A(c[1]) ->C(x1[.]{p}),A(c[.]) @ 'mod_rate' //AB modifies x1
'a.c' A(x[.],c[.]),C(x1{p}[.],x2[.]{u}) ->
      A(x[.],c[1]),C(x1{p}[.],x2[1]{u}) @ 'on_rate' //A binds C on x2
'mod␣x2' A(x[.],c[1]),C(x1{p},x2{u}[1]) ->
         A(x[.],c[.]),C(x1{p},x2{p}[.]) @ 'mod_rate' //A modifies x2

/* Variables */
%var: 'on_rate' 1.0E-3 // per molecule per second
%var: 'off_rate' 0.1 // per second
%var: 'mod_rate' 1 // per second
%obs: 'A' |A(x[.])|
%obs: 'AB' |A(x[x.B])|
%obs: 'Cuu' |C(x1{u},x2{u})|
%obs: 'Cpu' |C(x1{p},x2{u})|
%obs: 'Cpp' |C(x1{p},x2{p})|

%var: 'n_ab' 100
%obs: 'n_c' 1000
%var: 'C' |C()|

/* Initial conditions */
%init: 'n_ab' A(),B()
%init: 'n_c' C()
```

Figure S1: Comparison between Beacon Calculus (a) and Kappa (b) encodings of a simple protein-protein interaction network. The Kappa example is adapted from the manual, available at `https://kappalanguage.org/`. The result of one simulation of (a) using bcs is shown in Fig. S2a and the result of one simulation of (b) using the Kappa simulator is shown in Fig. S2b.

(a) (b)

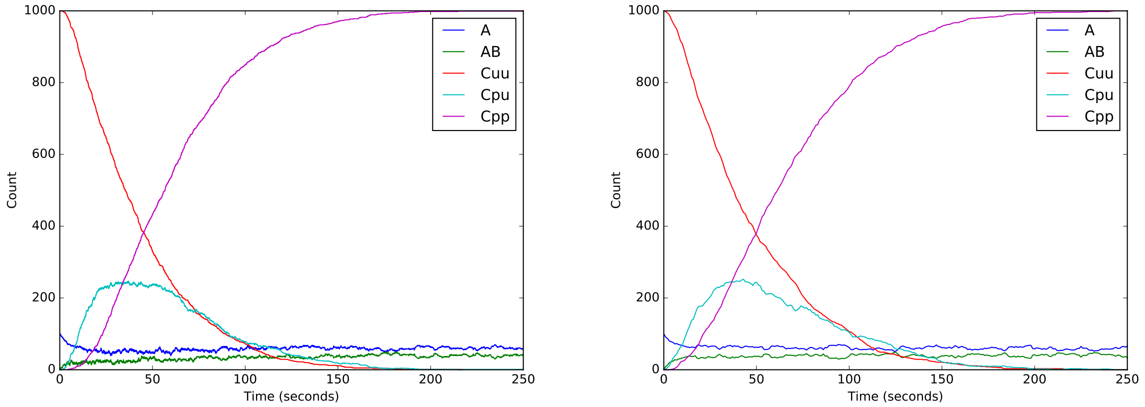


Figure S2: (a) The result of one simulation of the Beacon Calculus model in Fig. S1a simulated using bcs. (b) The result of one simulation of the Kappa model in Fig. S1b using the Kappa simulator.

#### S2.3 Comparison with PEPA (and its Variants)

The syntax of the Beacon Calculus is similar to Performance Evaluation Process Algebra (PEPA), using the notation of ordered pairs where each action is paired with the rate at which it is performed by a process. PEPA supports action prefix, choice, parallel composition of processes, and synchronised actions between processes [14]. The original formulation of the PEPA language does not contain features such as gates and parameters. These features (including using parameters in rate expressions) were added in the PEPAk extension of PEPA [6]. PEPAk also supports variable binding via handshakes in a similar fashion to the Beacon Calculus. One advantage of the Beacon Calculus over PEPA is that receiving messages over a set of values can make some models much more concise (see the examples in the methods section). In addition, PEPA models encode a finite continuous time Markov chain (CTMC) so that the system can be fully analysed. A consequence is that neither PEPA nor PEPAk support processes that can carry on as two parallel processes. Definitions such as

$$P_i = \{a, r\} . (P_{i+1} \parallel P_i) \tag{19}$$

are not allowed. The Beacon Calculus is intended for simulation, where mapping onto a finite CTMC is of limited use. Definitions of the type shown in Eqn. (19) are used in the DNA replication model, the DNA damage model, and the T-cell receptor model, so PEPA and PEPAk are not expressive enough to encode any of the three models in this paper. However, if the model does not require this feature, then PEPA can be adventageous: The representation of a model as a finite CTMC allows for the analysis of system properties (such as the asymptotic behaviour of the model, obtained by using the CTMC's stationary distribution). It should be noted that even if a model is finite, it can be computationally expensive to analyse the CTMC of a model with a finite but large number of states.

Bio-PEPA extends PEPA with a notion of compartments and the ability to cope with additional types of chemical kinetics (Hill kinetics, Michaelis-Menten kinetics) beyond mass action [5]. The Beacon Calculus can cope with these types of kinetics as well (depending on how the model is encoded) but in cases where a process models an individual molecule, reactions have an order of no more than two. Chemical reactions of order greater than two are, in general, difficult to encode as bimolecular stepwise reactions and often involves some knowledge or assumption about reaction intermediates. Bio-PEPA was designed to cope naturally with these cases and may be a better choice of tool than the Beacon Calculus if the system being modelled has more general kinetic laws or higher order reactions.

#### S2.4 Comparison with BioNetGen

BioNetGen is a rule-based language that allows users to specify reaction rates, components in the model (which are similar to processes in the Beacon Calculus), and reaction rules in order to generate a network of possible interactions [2, 12]. This network can be compiled to a set of ordinary differential equations (ODEs) or simulated using the Gillespie algorithm, thereby enabling the analysis of both deterministic and stochastic models. BioNetGen was designed for the analysis of models with high computational complexity, such as signalling networks where species in the system have a number of interactions or modifications that scales combinatorially.

BioNetGen was used by Dushek *et al.* in [11] in order to generate ordinary differential equations for the multisite phosphorylation model described in Fig. 3. The BioNetGen source code for this model was made available by the authors in the supplemental information of [11] and, as rendered in that document, is approximately 2.5 pages long consisting of 194 lines of code. As shown in Fig. 3, the corresponding Beacon Calculus model consists of 26 lines of code; it is considerably shorter and more concise. However, it should be noted that unlike the Beacon Calculus, BioNetGen models can be compiled into a set of ODEs which has an advantage for some applications: fitting ODEs to data is generally easier than fitting stochastic models, and ODEs facilitate useful analysis for some applications (such as phase plane analysis and bifurcation analysis).

#### S2.5 Comparison with PySB

PySB is a rule-based language that provides useful macros for common molecular processes in systems biology such as catalysis, pore assembly, synthesis, and reversible binding [16]. These are then compiled into BioNetGen or Kappa rules; these BioNetGen rules can then be converted into a system of ODEs and integrated using an external solver. A key advantage of PySB is that macros make code shorter and more reusable. In addition, PySB models are Python programs which can then be tracked using with version control software.

PySB uses BioNetGen to generate rules, which PySB then parses into ODEs which are integrated by an external tool (such as VODE). This flexibility can be of benefit, as the software maintains a wide range of compatibility with common tools such as NumPy, SciPy, and MATLAB. However, potential drawbacks are more complicated workflows as well as a number of dependencies that the user must seek out, install, and maintain. The bcs software is self-contained so that it parses the Beacon Calculus language and simulates the model without relying on external tools. An advantage of this approach is that users do not have to seek out and maintain dependencies, but a drawback is that it does not take advantage of other tools.

The use of pre-built macros for common model motifs in systems biology can represent an advantage over the Beacon Calculus, particularly if a model is mostly comprised of these motifs. As shown in the previous sections (particularly in Section S2.2 and S2.3) modelling large-scale regulatory or protein-protein interaction networks generally does not play to the Beacon Calculus's strengths and other tools such as PySB can be more appropriate. However, the DNA replication example in Fig. 1 of the main text describes the coordinated action of replication forks, which the Beacon Calculus is able to elegantly encode, particularly when the model is extended to include cooperative origin firing in Fig. 3. Macros of the type available in PySB may be less helpful when modelling this type of system. The Beacon Calculus finds its niche modelling systems that tend to be more "nonstandard" than conventional chemical reaction networks, such as the protein and damage levels within a population of dividing cells (Fig. 2) or the coordinated action of molecular machines (Fig. 1 and Fig. 4).

#### S2.6 Comparison with ML-Rules

ML-Rules is a rule-based language for systems biology that works as a plug-in for JAMES II [13, 17]. While ML-Rules is not a process calculus, it shares many features from process calculi such that it has the most in common with the Beacon Calculus out of the other tools mentioned in this section. In ML-Rules, species can have attributes (analogous to Beacon Calculus processes and parameters) and reaction rules can be written with patterns of these attributes to achieve similar behaviour to handshakes in the Beacon Calculus. Functions on these attributes can also be used in rate calculations for reactions, and ML-Rules permits predicates on these attribute values that determine whether a reaction is allowed to take place. One of the defining features of ML-Rules is the compartmentalisation of species such that they can be nested.

In addition to the conceptual difference between a rule-based language and a process calculus, the main difference brought by the Beacon Calculus is the option for processes to interact with a global database via beacons. One example of when this is useful for biological applications is a kinesin motor stepping down a microtubule where it can blocked by diffusible Tau proteins (Section S5). In the Beacon Calculus, the Tau protein can be modelled as a process that maintains an active beacon with its location, and kinesin motors can use a beacon check to determine whether a Tau protein is nearby and remain blocked until the Tau protein diffuses away. Similar behaviour can occur in DNA replication, as DNA replication forks can be blocked by DNA binding proteins and restarted once the obstruction is removed. Implementing this blocking behaviour is less straightforward to implement in ML-Rules. However, a key feature of ML-Rules is compartmentalisation, and while none of the biological applications in this paper relied on compartmentalisation, ML-Rules may be a more appropriate choice over the Beacon Calculus for applications where this is important.

##### S2.7 Comparison with Simmune

Simmune is a rule-based language tailored to immunology, though it can be used across other disciplines as well [1, 23]. It provides a graphical user interface where users can specify molecules, their individual binding sites, and the state of the molecules in order to model their interactions. Molecules can also have ownership of components such that, for example, interacting components can be specified on the exterior or interior of a membrane. This is particularly important for immunology applications, as the "molecules" will often be cells where the ligands can interact with receptors that are external to the cell, producing some intra-cellular state change. Critically, Simmune can also simulate spatial interactions and constraints so to produce realistic models of, for example, cell-to-cell contact formation.

The graphical user interface of Simmune makes modelling more accessible, particularly to nonspecialists. For applications where there are combinatorial interactions between multiple receptors or sites on different molecules, and particularly when the spatial aspect of these interactions is important, Simmune is clearly the stronger choice over the Beacon Calculus. For the three applications shown in the main text, these aspects were not particularly important, and the Beacon Calculus was able to encode these biologically meaningful scenarios elegantly and easily. Therefore, the Beacon Calculus and Simmune occupy different niches and can be seen as complementary.

#### S3 DNA Replication Model System Line

For the DNA replication models in Fig. 1 and Fig. 4, the system line of the DNA replication model has been truncated for clarity. S. cerevisiae chromosome II, on which these models are based, is comprised of approximately 33 known origins of replication. Each of these origins, along with their chromosomal coordinate, licensing probability, and mean firing time, need to be specified on the system line as an initial condition of the model. These parameters, which are available from [9], are easily parsed into the system line below using a scripting language.

```
L = 813; //length of S. cerevisiae chrII in kilobases
fast = 100000; //fast rate
v = 1.4; //fork velocity in kilobases per minute

//process definitions
ORI[i,fire,q] = {licensed,q*fast}.{~chr?[i],fire}.(FL[i]||FR[i])
                + {nlicensed,fast*(1-q)};
FR[i] = {chr![i],fast}.[i < L] -> {~chr?[i+1],v}.FR[i+1];
FL[i] = {chr![i],fast}.[i > 0] -> {~chr?[i-1],v}.FL[i-1];

//initial processes in the system
Ori[1,0.025,0.522] || Ori[6,0.201,0.097] || Ori[29,0.036,0.745] ||
Ori[63,0.052,0.646] || Ori[93,0.045,0.354] || Ori[143,0.037,0.76] ||
Ori[170,0.031,0.724] || Ori[177,0.031,0.127] || Ori[198,0.040,0.918] ||
Ori[209,0.031,0.869] || Ori[237,0.064,0.793] || Ori[255,0.040,0.916] ||
Ori[283,0.043,0.147] || Ori[326,0.036,0.916] || Ori[378,0.037,0.592] ||
Ori[389,0.055,0.267] || Ori[407,0.028,0.941] || Ori[417,0.039,0.72] ||
Ori[441,0.105,0.047] || Ori[486,0.050,0.792] || Ori[517,0.044,0.234] ||
Ori[539,0.054,0.379] || Ori[591,0.053,0.497] || Ori[612,0.030,0.982] ||
Ori[622,0.047,0.948] || Ori[632,0.034,0.747] || Ori[676,0.045,0.15] ||
Ori[706,0.048,0.534] || Ori[720,0.050,0.286] || Ori[741,0.035,0.861] ||
Ori[757,0.032,0.655] || Ori[774,0.036,0.689] || Ori[794,0.025,0.284] ||
Ori[802,0.035,0.817];
```

Figure S3: DNA replication model written in the Beacon Calculus with the full system line specified for *S. cerevisiae chromosome II.*

#### S4 Benchmarks

To give users a practical sense of how long it takes to simulate Beacon Calculus models using the bcs software, this section provides benchmarks for the examples simulated in this paper. All benchmarks were done using an Intel(R) Core(TM) i7-9700 CPU @ 3.00GHz CPU on a single thread. While we do not use multithreading in these benchmarks, the bcs software can easily run simulations in parallel by running a separate simulation on each thread which can lead to much faster run times than those shown below as most modern computers have at least four cores. Note that bcs simulations are not memory-intensive and system memory should not be an issue for simulations.

In the table below, each row specifies the model, the number of simulations run, simulation termination conditions, the results of five trials, and the average run time of these five trials. All times are specified in seconds. For the DNA damage model, benchmarks are given for the model with both low and high rates of DNA damage.

| **Model** | **Sim.** | **Termination** | **Trial 1** | **Trial 2** | **Trial 3** | **Trial 4** | **Trial 5** | **Average** |
|---|---|---|---|---|---|---|---|---|
| DNA Replication | 500 | deadlock | 27.726 | 27.852 | 27.096 | 27.721 | 28.026 | 27.684 |
| DNA Damage (Low) | 25 | 250 minutes | 1.863 | 2.411 | 1.619 | 4.063 | 2.45 | 2.481 |
| DNA Damage (High) | 25 | 250 minutes | 89.534 | 63.705 | 48.883 | 101.251 | 49.582 | 70.591 |
| Multisite Phos. | 50 | 5000 seconds | 203.012 | 205.798 | 2011.417 | 205.685 | 204.859 | 206.154 |

#### S5 Additional Example: Kinesin Stepping Down a Microtubule

This section provides an alternative way to introducing the Beacon Calculus language, providing an additional introductory example to complement those in the Language Overview subsection of the main text. In particular, it describes kinesin motors moving down a microtubule (Fig. S4). While this model provides a framework that could be further developed to provide biological insight, this example is intended for demonstration and the parameter values chosen do not necessarily reflect biologically realistic values.

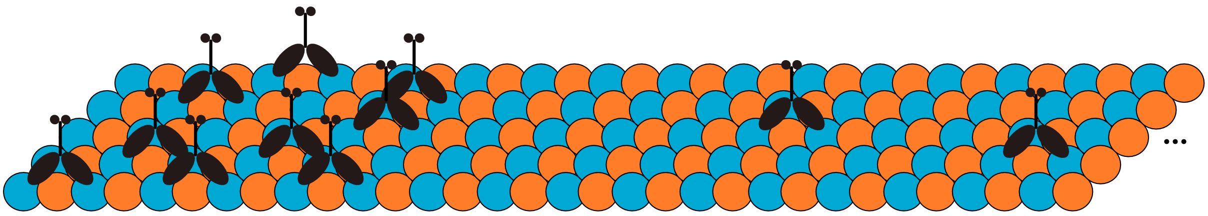

Figure S4: **Molecular crowding of kinesin stepping down a microtubule.** Kinesin (black) stepping down dimers of tubulin (blue, orange) in a microtubule. Each kinesin motor is a process `K` that can interact with motors around it via beacons and handshakes.

The following is a very simple Beacon Calculus model that describes the stepping of the motor protein kinesin down a microtubule:

```
1 r = 1; //define the rate for walking
2
3 //process definition
4 K[i] = {walk,r}.K[i+1];
5
6 //give the initial state of the system
7 K[1];
```

In this model, a kinesin process `K` has a position given by its parameter `i`. It starts at position `i=1` (Line 7) and then walks at rate `r`. Every time it takes a step, it recurses with the value of its parameter increased by one. Suppose instead that kinesin should accelerate forward until location `i=5` and then stop. The parameter `i` can be used to scale the static constant `r` so that the rate of walking increases as the motor gets closer to `i=5`. This can be expressed as follows:

```
r = 1; //define the rate for walking

//process definition
K[i] = [i<5] -> {walk,r*i}.K[i+1];

//give the initial state of the system
K[1];
```

Once `i=5`, the condition specified in the gate no longer holds and kinesin cannot perform the action `walk`. When a process can no longer perform any actions, it is said to be deadlocked and is removed from the system. If all processes in the system are deadlocked, the simulation stops.

The following example includes two kinesin motors stepping on the same microtubule.

```
1  r = 1; //define the rate for walking
2  rr = 2; //handshake receive rate
3  rs = 3.5; //handshake send rate
4  L = 100; //microtubule length
5
6  //process definition
7  K[i] = {@bump?[i+1],rr}.K[i]
8       + {@bump![i],rs}.K[i]
9       + [i<L] -> {walk,r}.K[i+1];
10
11 //initial state of the system
12 K[5] || K[8];
```

This model has two kinesin motors: one starts at `i=5` and the other starts at `i=8`. They can communicate over channel `bump` (Lines 7-8). If they step such that one kinesin is directly ahead of the other (if they were respectively at positions `i=10` and `i=11`, for example) then they can handshake over channel `bump` at rate `rs*rr`. If they do, both kinesin motors remain where they are. This handshake, when it is possible, competes with each motor's ability to step independently (Line 9). Therefore, the handshake has the effect of the motors impeding each other's progress when they are nearby.

If a handshake receive can accept multiple values, the receiving process can bind the value it receives to a variable for later use. The process may, for instance, use this value in a rate expression or as a parameter. The binding variable can be used in the rate expression to indicate how different values can be received at different rates; it can bias which value in the set is received. For example, suppose it is more likely that two kinesin motors impede each other as they get closer to one another. The two definitions for kinesin below, `K1` and `K2`, are equivalent.

```
r = 1; //define the rate for walking
rs = 3.5; //handshake send rate
L = 100; //microtubule length

//process definition
K1[i] = {@bump?[i+1..i+5](x),1/abs(i-x)}.K1[i]
      + {@bump![i],rs}.K1[i]
      + [i<L] -> {walk,r}.K1[i+1];
K2[i] = {@bump?[i+1],1}.K2[i]
      + {@bump?[i+2],1/2}.K2[i]
      + {@bump?[i+3],1/3}.K2[i]
      + {@bump?[i+4],1/4}.K2[i]
      + {@bump?[i+5],1/5}.K2[i]
      + {@bump![i],rs}.K2[i]
      + [i<L] -> {walk,r}.K2[i+1];

//initial state of the system
K1[5] || K2[8];
```

Tau is a neuronal microtubule associated protein that diffuses up and down microtubules [15] and increases the rate of kinesin unbinding [4, 20]. In the following example, a tau protein diffuses on the microtubule and keeps a beacon active at its current position. Kinesin has a chance to unbind from the microtubule when it is at the same position as tau.

```
1  f = 10000; //fast rate
2  r = 1; //kinesin stepping rate
3  rr = 2; //handshake receive rate
4  rs = 3.5; //handshake send rate
5  d = 5; //tau diffusion rate
6  L = 100; //microtubule length
7
8  //process definition
9  tau[i] = [i>0] -> {left,d}.{block#[i],f}.{block![i-1],f}.tau[i-1]
10        + [i<L] -> {right,d}.{block#[i],f}.{block![i+1],f}.tau[i+1];
11
12 K[i] = {@bump?[i+1],rr}.K[i]
13      + {@bump![i],rs}.K[i]
14      + [i<L] -> {walk,r}.K[i+1]
15      + {block?[i],r};
16
17 //initial state of the system
18 K[5] || K[8] || tau[10];
```

In this model, tau diffuses left (Line 9) or right (Line 10) along the microtubule. Both the `left` and `right` actions have the same rate `d`, so the diffusion is unbiased. Tau keeps a beacon active on channel `block` that transmits its position. When it moves left or right, it kills the beacon transmitting its current location and launches a beacon transmitting its new location. These beacon actions should happen immediately so that their rates do not impact the rate of left-right diffusion; they are done at a fast rate (Line 1) relative to other action rates in the system. If there is a beacon active on channel `block` transmitting the same value as kinesin's location, there is a chance for kinesin to receive the beacon (Line 15) and deadlock.
